# Programmable Aggregation of Artificial Cells with DNA Signals

**DOI:** 10.1101/2020.10.18.344432

**Authors:** Hengming Qiu, Feiran Li, Yancheng Du, Ruixin Li, Ji Yeon Hyun, Sei Young Lee, Jong Hyun Choi

## Abstract

Cell aggregation is a complex behavior, which is closely related to the viability, differentiation, and migration of cells. An effort to create synthetic analogs could lead to considerable advances in cell physiology and biophysics. Rendering and modulating such a dynamic artificial cell system require mechanisms for receiving, transducing, and transmitting intercellular signals, yet effective tools are limited at present. Here we construct synthetic cells from engineered lipids and show their programmable aggregation behaviors using DNA oligonucleotides as a signaling molecule. The artificial cells have transmembrane channels made of DNA origami that are used to recognize and process intercellular signals. We demonstrate that multiple small vesicles aggregate onto a giant vesicle after a transduction of external DNA signals by an intracellular enzyme, and that the small vesicles dissociate when receiving ‘release’ signals. This work provides new possibilities for building synthetic protocells capable of chemical communication and coordination.

---

Cell aggregation is an important phenomenon in biology, where cells cluster together upon external cues in a certain environment. Studying the intercellular signaling and cell-cell interactions during the process can thus contribute substantially to the understanding of cell differentiation, migration, and viability.^1–3^ It also holds great potential in enhancing tissue engineering research as cell interactions are an essential part of tissue building.^4^ Tissue engineering requires not only scaffold matrices that provide structural supports, but also cellular signals to generate growth factors.^5^ Given the importance, considerable research efforts were made on this topic, including signaling mechanisms and associated molecules, the kinetics and dynamics of cell aggregation, and the related diseases.^6,7^ With the complexity of cellular environments, however, it is still a significant challenge to fully understand the aggregation behaviors and actively control the processes. Therefore, a simplified synthetic system capable of cell-to-cell communication and coordination would help achieve a deeper understanding and precise control of cell interactions and behaviors.^8^ Engineered lipid vesicles may serve as such a versatile platform with cell-like structures and properties, thus resembling some characteristics of biological cells.^9^

Over the past several years, DNA nanotechnology has been explored to construct and control synthetic cells. DNA self-assembly can produce complex architectures with sub-nanometer precision,^10–12^ dynamic nanostructures such as switches and motors,^13,14^ and computing devices.^15^ The key advantage is DNA’s excellent programmability and structural predictability. For example, DNA assemblies were used to stabilize vesicle structures and control their shapes.^16,17^ DNA base-pairing was implemented for programmable vesicle fusion.^18,19^ Recently, DNA origami was used to mimic the shape and function of naturally occurring membrane protein channels.^20–22^ They were incorporated into lipid bilayer membranes and served as pores for biomolecular transport in and out of vesicles.^23^ The kinetics of tranport process through DNA origami pores was measured with dye molecules.^24^ The geometry and chemical functionality of this novel class of artificial nanopores can be rationally designed using computer-aided molecular engineering tools that are available for DNA nanotechnology.^25–28^ Protein or peptide membrane pores also offer molecularly defined dimensions; however, their geometry cannot be modified as easily as for DNA nanostructures, and their chemical functionalization typically is more cumbersome.^29^ These features make DNA-based membrane channels highly promising biomolecular devices for applications in single-molecule biosensing and drug delivery, or as components for artificial cells.^30^

In this study, we demonstrate programmable communication and coordination between artificial cells by using transmembrane pores made of DNA origami. The engineered vesicles interact with each other and aggregate controllably and reversibly, as biological cells do. We develop new signaling mechanisms such that the artificial cells follow biochemical instructions for clustering behaviors. A set of external biomolecular signals are recognized and transported into the vesicles, where they are transduced to another form of signals that move out of the cells through the origami membrane channels, thereby initiating cell aggregation. We use DNA oligonucleotides as singling molecules to trigger association and dissociation of vesicles, which is monitored by fluorescence microscopy. This system mimics biological cell behaviors and provides useful tools for cell biology research and bottom-up synthetic biology.

Figure 1 presents the scheme of our experiment. Two types of DNA-decorated vesicles are prepared. Giant unilamellar vesicles (GUVs) were assembled by an inverted emulsion method, while small unilamellar vesicles (SUVs) were synthesized with a dehydration-rehydration method (see SI for experimental details).^31,32^ The GUV contains tubular DNA origami pores that are designed to have a diameter of ~32 nm and a length of ~60 nm (Figure S1). Each tubule is functionalized with cholesterol moieties at the half length to insert the origami in the GUV during the assembly and stabilize it in the hydrophobic environment of the lipid membrane. The DNA pores are initially closed with rectangular DNA origami caps by using two sets of staple extensions that are complementary to each other. A set of staples on the tubular pore have a 16-nucleotide (16-nt) extension, while the rectangular cap includes staples with a 24-nt extension (shown in pink and green in Figure S1, respectively). These staples are used to attach the rim of the DNA tubule with the 60-nm×100-nm rectangular tile. The atomic force microscopy (AFM) images indicate more than 90% of the tubular structures are linked with flat origami (Figure S2). Giant vesicles are immobilized on a glass coverslip via biotin-streptavidin conjugation (Figure 1a). The glass surface is passivated with biotin functionalized bovine serum albumin (BSA), and then streptavidin is introduced. The GUV containing biotinylated phospholipids thus binds to the surface and remains there during the experiments.

**Figure 1.**
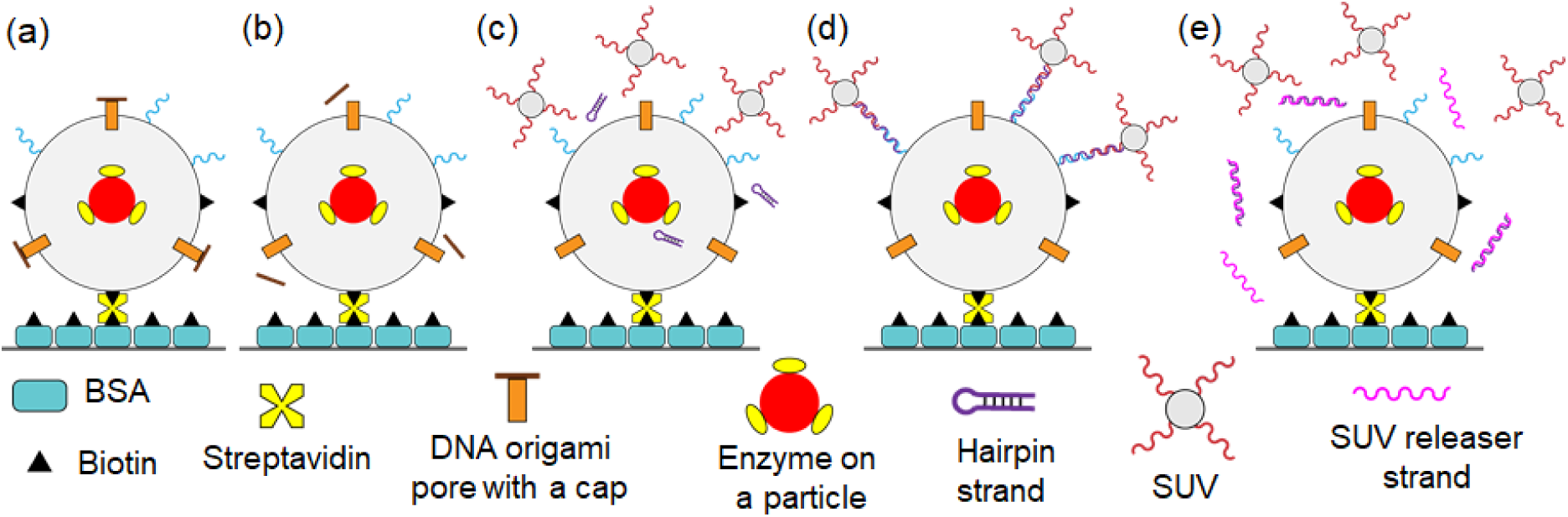
Scheme for reversible cell aggregation programmed by DNA signals. (a) A giant vesicle is immobilized on a BSA modified glass surface by biotin-streptavidin conjugation. The GUV has DNA strands (shown in blue), includes transmembrane channels made of DNA origami tubules (orange), and encapsulates Exo III on a polystyrene particle (yellow-red). (b) The pores are initially closed with flat origami caps which can be removed by ‘cap-releaser’ strands via toehold-mediated strand displacement. (c) An external DNA hairpin signal can enter the vesicle through the membrane channels where it is transduced into another form of signal via enzymatic reaction by Exo III. The enzyme digests a part of the hairpin from the 3’ end, thereby exposing the complementary domains for the strands on the vesicles (both blue and red). (d) The processed signaling oligonucleotides will pass through the origami pores and triggers aggregation of multiple small vesicles on the GUV. (e) Another DNA signal, ‘SUV releaser’ strands shown in pink, will dissociate the SUVs bound on the giant vesicle, by removing the linker strands. This association and dissociation behavior can be programmed with DNA signals and repeatedly indefinitely in theory.

The DNA pores, initially closed with flat DNA origami caps, can be opened using a set of ‘cap releaser’ strands. The staple extensions on the rectangular origami tile has an 8-nt single-stranded overhang at the 3’ end such that fully complementary cap releasers can remove them via toehold-mediated strand displacement,^33^ as illustrated in Figure 1b. A hairpin, shown in purple, is used as a signal for cell-cell interactions. This strand has two domains; one is complementary to the strand on the giant vesicle (shown in blue), and the other can hybridize with the DNA on the small vesicles (red). However, these domains are initially shielded within the hairpin structure, thus it cannot bind to the DNA strands on the vesicles. With the cap removed, the hairpin can enter the GUV via diffusion, where it can be opened by enzymatic cleavage (Figure 1c). Exo III enzymes are functionalized on a polystyrene particle with a diameter of ~200 nm, which are encapsulated in the giant vesicle. The cleaved DNA signal can then diffuse out of the GUV and connect the strands on both giant and small vesicles together, resulting in SUV aggregations on the GUV (Figure 1d). Rhodamine B is included on the membrane of the small vesicles, which is used to image the SUV aggregation. The small vesicles will disaggregate from the giant vesicle upon introduction of ‘SUV releaser’ strands via strand displacement as the releaser strands are fully complementary to the SUV-linkers (Figure 1e).

Figure 2 confirms the giant vesicles with DNA origami pores. The synthesized GUVs have a range of diameters from 5 to 25 µm. Here we decorated the origami tubules with Cy5 dye for optical imaging (Figure S1). The giant vesicles were immobilized on the bottom surface of the microfluidic channel, and the imaging was performed under an inverted microscope using a 63x objective lens. In the fluorescence image, a distinct ring in pseudo red color is seen around the vesicle compared with the brightfield image (Figure 2a), confirming the presence of DNA origami pores. In contrast, no fluorescence was observed from the GUV when cholesterol was not used in DNA origami (Figure 2b). This indicates that DNA tubules are not embedded in the lipid bilayer, as anticipated. We then tested molecular diffusion through open transmembrane channels on the GUV containing origami tubules (without caps). Here, green fluorescent protein (GFP) and Cy5-labelled 60-nt DNA were used in the measurement. The fluorescent molecules were supplied to the giant vesicles in tris-acetate-ethylenediaminetetraacetic acid (EDTA) buffer containing 10 mM MgCl2 (termed TAEM buffer). The fluorescence intensity inside the GUVs increased shortly after the inflow, as shown in Figure 2c and 2e. Without origami pores, however, fluorophores cannot enter the vesicles, thus no fluorescence was detected (Figure 2d and 2f).

**Figure 2.**
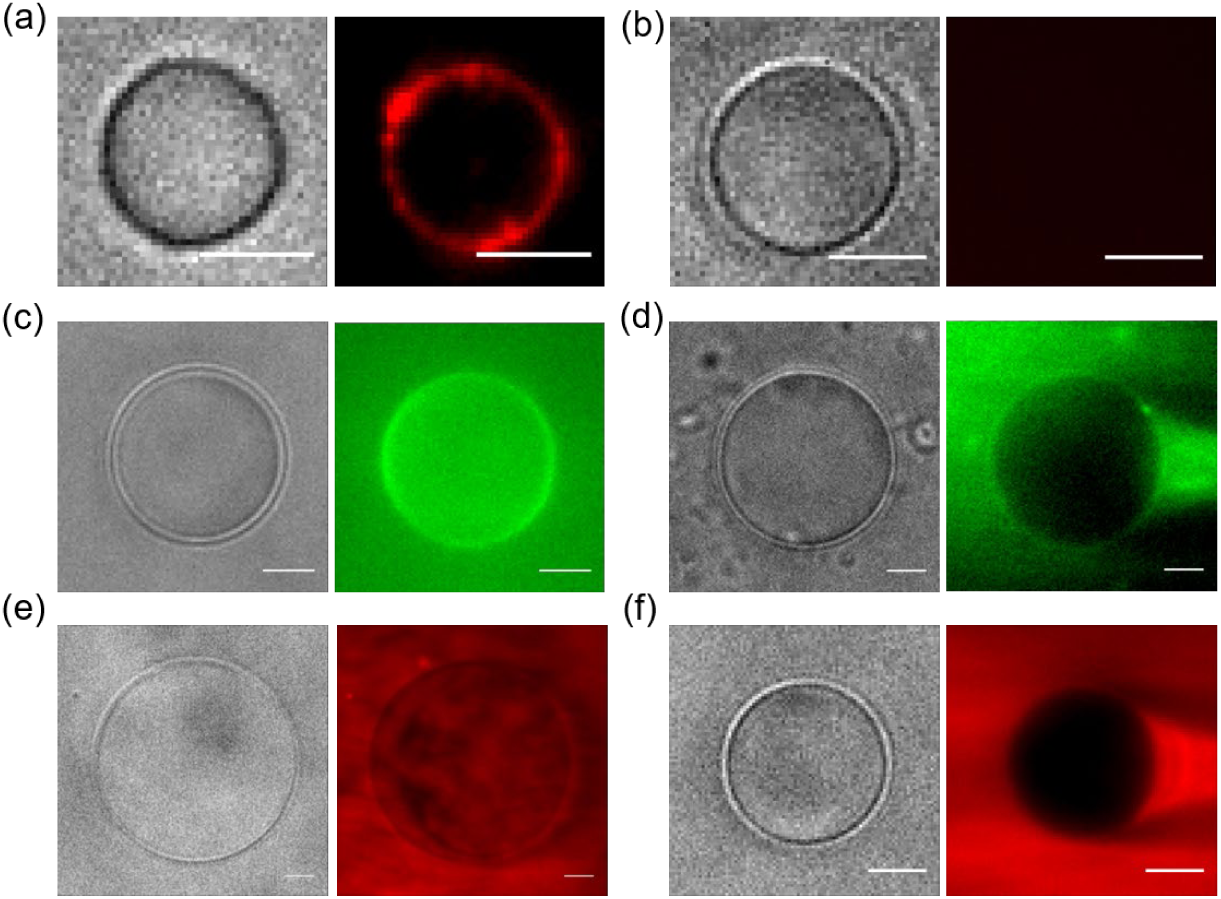
Brightfield (left) and fluorescence (right) images of giant vesicles. In fluorescence images, GFP is shown in green color and Cy5 is represented in red. (a) Cholesterol modified DNA tubules can be incorporated into vesicle membrane. The Cy5 functionalized origami shows a distinct, ring-shaped fluorescence pattern around the vesicle boundary. (b) DNA origami without cholesterol moieties cannot insert in the membrane, thus no fluorescence is observed. (c) GFP can penetrate into the vesicle via transmembrane channels made of tubular origami. (d) Without DNA tubules, GFP cannot diffuse into the vesicle. (e) Cy5-DNA can enter the vesicle with DNA pores. (f) The fluorophore labelled DNA cannot move into the GUV, when no origami pores are present on the vesicle surface. Note that the origami tubules in (c) and (e) are not functionalized with Cy5 dyes. Scale bars are 5 µm.

To characterize the outflux through the membrane channels, we measured the kinetics of GFP and Cy5-DNA from the giant vesicles (Figure 3). The fluorescent molecules were initially encapsulated inside the GUVs containing the DNA pores with the flat origami caps on. The fluorescence intensity does not change initially, suggesting that no significant leakage of dye molecules with the origami pores closed. When the cap releasers were supplied to the channel (indicated by the orange arrows in Figure 3b and 3d), a drastic decrease of fluorescence intensity inside the vesicle was observed within a few minutes. The fluorescence dropped until it reaches around 40% of the initial intensity. To show the drastic changes, we overlaid the fluorescence images (shown in pink) at the beginning and end of the measurements with the brightfield images (Figure 3a, 3c). It is worth noting that the outfluxes of GFP and Cy5-DNA are similar and that a fraction of fluorescent molecules remains inside the vesicle after a long period of time. The fluorescence decrease was curve-fitted with a single-exponential function, yielding a time constant of ~3 min for GFP and ~8 min for Cy5-DNA (see SI for details). These timescales agree well with the previously reported value (~15 min) using 40 kDa dextran with fluorescent moiety by Thomsen *et al*.^24^

**Figure 3.**
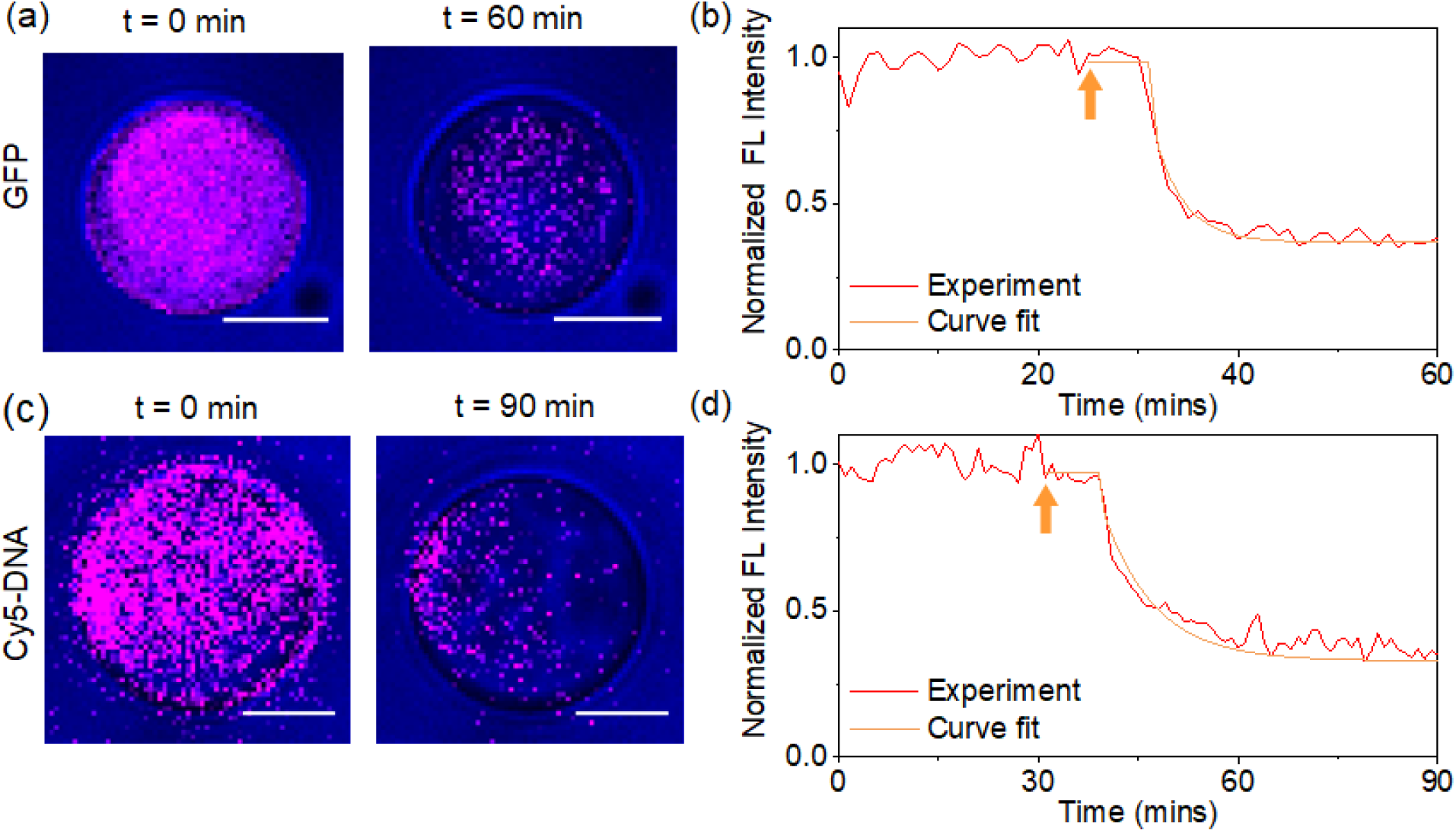
Kinetic measurement of molecular outflux through origami pores from giant vesicles immobilized on the glass coverslip surface. (a) Fluorescence images of GFP molecules (shown in pink color) overlaid with brightfield images of a vesicle at time *t* = 0 and 60 min. The molecules, initially encapsulated in the GUV, diffuse out of the vesicle after the pores are opened. (b) The fluorescence intensity inside the vesicle decreases, shortly after the cap-releaser strands were introduced into the microfluidic imaging chamber (indicated by the orange arrow). The exponential curve-fit (orange line) suggests a time constant of ~3 min. (c) Fluorescence images of Cy5-DNA inside a giant vesicle overlaid with brightfield images at time *t* = 0 and 90 min. (d) The fluorescence diminishes after adding the cap releasers (orange arrow) with a ~8-min time constant. Scale bars are 5 µm.

Next, we examined the DNA-programmed reversible clustering of vesicles. To test the effectiveness of the SUV linkers and releasers, the origami pores and the Exo-III modified particles were not included in the large vesicle, while the SUVs and GUV have unique strands shown in red and blue in Figure 1. We first immobilized a large vesicle on the imaging surface (state (i), Figure 4a). Then, 36-nt SUV linkers with complementary domains to the strands on the small and large vesicles were supplied along with SUVs to the microfluidic channel (indicated by the red arrow). The final concentrations were about 1011/ml for the SUVs and 10 nM for the SUV linkers. After 10 min incubation, the TAEM buffer was supplied to wash away unbound small vesicles and excess linker strands. Distinct fluorescence is observed around the large vesicle boundary (state (ii)), indicating that the small vesicles indeed aggregate on the GUV surface as designed. Note that the fluorescence intensity originate from the SUVs as the giant vesicle does not include any fluorophores. We then supplied the SUV releaser strands to the imaging chamber, indicated by the black arrow. The invading strands first engage with the SUV linkers via 6-nt toehold and bind with them fully, removing the linkers from both the small and large vesicles. As a result, the SUVs dissociate from the GUV and the fluorescence intensity drops, as seen in state (iii). The aggregation and disaggregation of the small vesicles on the GUV were repeated by adding the linkers and releasers in series. The fluorescence intensity changed accordingly as shown in Figure 4a. In theory, such a reversible aggregation may be cycled indefinitely. In a control experiment, we replaced the SUV linkers with strands of random sequence (indicated by the blue arrows in Figure 4b) which should not base-pair with the strands on the vesicles. Indeed, vesicle aggregation was not triggered, and no significant changes in the fluorescence intensity were observed as shown in Figure 4b. This confirms that only designed DNA signals can lead to the aggregation behaviors.

**Figure 4.**
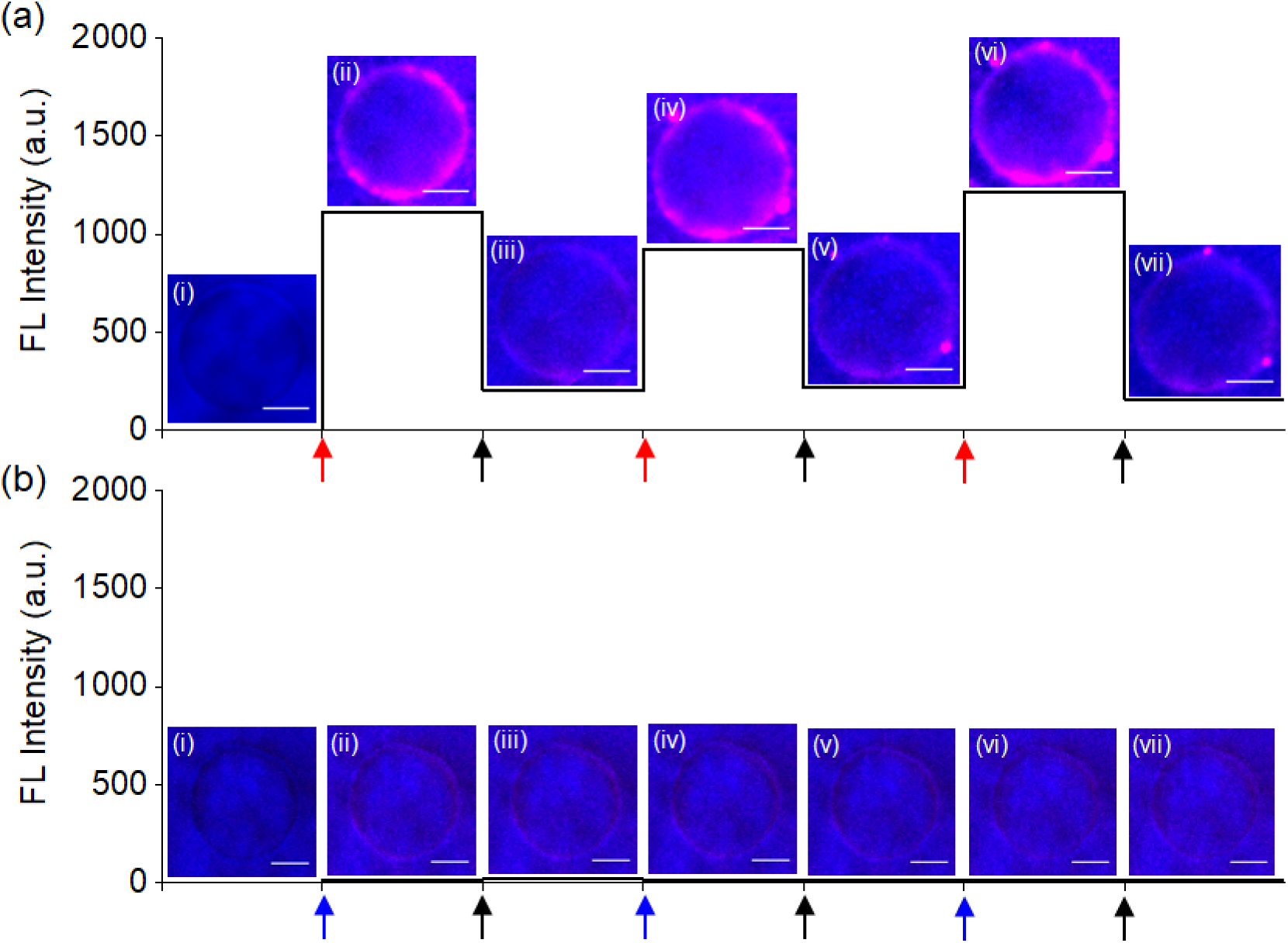
Reversible vesicle aggregation with DNA signals. (a) Fluorescence intensity of the small vesicles around the GUV on each experimental step. Corresponding images of fluorescence overlaid on brightfield images are also presented. The SUVs have oligonucleotides (red strands in Figure 1) and rhodamine B dyes for imaging. The giant vesicle contains DNA strands on its surface (blue strands in Figure 1), but does not include DNA origami pores nor polystyrene particles. (i) A giant vesicle is immobilized on the microfluidic imaging chamber via biotin-streptavidin conjugation. (ii) Small vesicles and 36-nt SUV-linker DNA are introduced (indicated by the red arrow) which will trigger SUV clustering on the giant vesicle via base-pairing of the linkers with the strands on both giant and small vesicles. As a result, the fluorescence intensity increases drastically. (iii) A set of SUV releaser strands are supplied, represented by the black arrow. The signaling oligonucleotides will first bind with the 6-nt toehold and hybridize fully with the SUV-linkers, thus removing the linkers from the vesicles. The small vesicles will thus dissociate from the GUV, and fluorescence intensity drops drastically. (iv)-(vii). The reversible vesicle aggregation can be repeated with DNA signals (SUV linkers and releasers). (b) Control experiment, using a random DNA sequence instead of the SUV linkers (represented by the blue arrows). As anticipated, the small vesicles are not bound on the GUV, thus no significant change in the fluorescence is observed. Scale bars are 5 µm in all images.

Finally, we demonstrated the recognition and transduction of DNA signals through membrane pores for programmable vesicle aggregation. Here we used the 56-nt hairpin that contains complementary domains for the strands on the vesicles (*i.e*., SUV linker sequence). As discussed in Figure 1, the domains are initially shielded so that it cannot induce aggregation directly (Figure S9). This hairpin signal may be transduced by Exo III to expose the SUV-linker domain via enzymatic digestion. The Exo III-modified, fluorescein-labelled polystyrene particles were encapsulated in the large vesicle during the assembly. The enzyme-particles are too large to pass through the DNA origami pores, thus they float randomly inside the GUV, shown as yellow dots in the fluorescence image in Figure 5a. The signal recognition and transduction were demonstrated using this large vesicle with the DNA pores initially closed by flat origami caps (state (i)). We first opened the DNA pores with the cap releasers, as indicated by the orange arrow (state (ii)). The hairpins can then enter the vesicle via membrane channels and interact with Exo III enzymes such that the shielding domain will be cut off and the 36-nt SUV-linker signal will be exposed. When the SUV linkers move out of the GUV, they can bind with the oligonucleotides on the small and large vesicles. As a result, the SUVs aggregate on the GUV, showing the circular fluorescence pattern around the large vesicle (state (iii)). The clustered, small vesicles were then dissociated (state (iv)), when the SUV releasers were introduced (indicated by black arrows), similar to what is seen in Figure 4. This programmable clustering was repeated twice, which may be continuously cycled as long as the chemical wastes are removed and new DNA signals are provided. As a control experiment, we replaced the hairpin strand with a random sequence. The small vesicles did not aggregate on the GUV with no significant fluorescence change observed, as expected (Figure 5b). It is worth mentioning that the shape of immobilized vesicles may change slightly over time as seen in Figure 5a as well as Figures S6 and S8. It is also notable that with the release signals the fluorescence becomes very weak, yet it does not drop to 0 (Figure 5a). This may be attributed to some nonspecific binding of small vesicles. Nonetheless, our experiment clearly demonstrates the effectiveness of DNA signals in programming aggregation behaviors of artificial cells.

**Figure 5.**
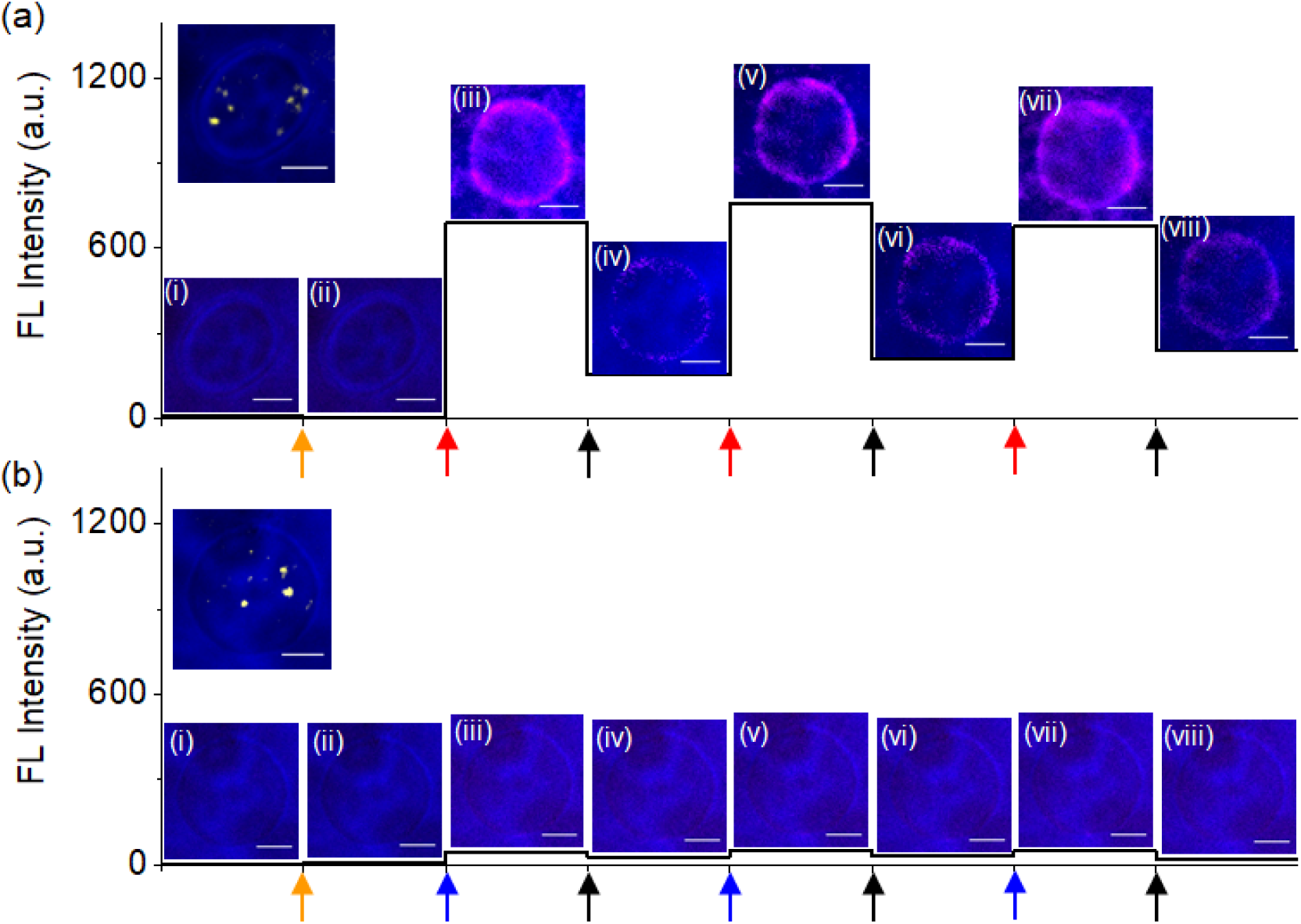
(a) Dynamic aggregation behavior of artificial cells programmed by DNA signals. To receive and transmit signals, tubular origami pores are included in the giant vesicle which also encapsulates Exo-III modified polystyrene particles. The 200-nm-diameter particles, shown as yellow dots in the upper left image, randomly float inside the GUV. (i) A giant vesicle is immobilized on the imaging chamber. (ii) A set of cap releaser strands are supplied to remove the flat origami caps (orange arrow). (iii) The hairpin strands and small vesicles are introduced to the channel, as indicated by the red arrow. The hairpins penetrate into the vesicle through the transmembrane origami channels and are partly digested by Exo III, exposing the SUV linker domain. The transduced oligonucleotides diffuse out through the pores and bind with the strands on the small and large vesicles. As a result, the small vesicles cluster around the GUV surface, which is evident with a circular fluorescent image overlaid on the brightfield image of the giant vesicle. (iv) When the SUV releasers are introduced (represented by the black arrow), they remove the SUV linkers via toehold-mediated strand displacement. The small vesicles then dissociate from the GUV, and the fluorescence intensity drops drastically. (v)-(viii) The DNA programmable clustering behavior can be repeated multiple times. (b) Control experiment using a random sequence instead of the hairpin. (i)-(ii) The DNA pores are opened by removing the caps (indicated by the orange arrow). (iii) The oligonucleotides of random sequence may enter the giant vesicle (blue arrow), but they cannot trigger the aggregation of small vesicles on the GUV. Thus, no significant changes in the fluorescence are observed (iii). (iv) The addition of SUV releasers will not result in any changes in the behavior of the vesicles. Scale bars are 5 µm in all images.

In summary, we have demonstrated chemical communication and cooperative behavior of synthetic cells using DNA signals. The key hairpin signal was effective for the purpose, when and only when it was allowed to enter the cell through membrane channels and interact with enzymes. The DNA origami based transmembrane channels were critical for receiving, transducing, and transmitting biochemical signals. The interaction between vesicles following DNA signals shows great potential for more complex multi-cellular behaviors. This study will help construct synthetic cells as a surrogate model system for cell biology and neuroscience research. For example, an artificial neuronal network could be constructed to study mechanisms for transferring and receiving neurotransmitters. Such a platform could elucidate propagation pathways of sensation such as pain, which has been traditionally difficult to study. Overall, we envision that the growing library of DNA nanotechnology tools will open new opportunities for both fundamental sciences and novel biotechnology applications.

## ACKNOWLEGEMENT

This work was funded by the U.S. Office of Naval Research. J.H.C. is grateful for partial support from the U.S. National Science Foundation and Department of Energy. S.Y.L. acknowledges the support from the Basic Science Research Program through the National Research Foundation of Korea (NRF) funded by the Ministry of Education (NRF-2016R1D1A1A02937019), Republic of Korea.

