## Supplementary Information for "Programmable Aggregation of Artificial Cells with DNA Signals"

**Table of content**

1. Materials
   - DNA sequence information
   - Origami design
   - Lipid information
2. Sample Preparation Methods
   - Conjugation of DNA to lipid molecules
   - Polystyrene particles with Exo III
   - Synthesis of giant vesicles
   - Small vesicles with DNA strands
   - Assembly of DNA origami structures
3. Experimental Setup
   - Flow channel assembly
   - Surface passivation
   - Imaging system
4. Characterization
   - AFM images of tubular and rectangular DNA origami
   - Additional kinetic measurement
   - Exo III activity inside a giant vesicle
   - Immoblized giant vesicle shape change over time
   - Reversible aggregation behavior using DNA signals
5. Kinetics
6. References

### 1. Materials

#### DNA sequence information

All the sequences were purchased from Integrated DNA Technologies and used without further purification.

**Table S1.** Sequences of DNA linker, releaser, and modified strands. The SUV linker can bind with both SUV and GUV strands, thereby connecting SUVs on a GUV. The hairpin strand includes the sequence of SUV linker which may be exposed after digestion of ACA TCT AAC AAC CAA ACC AT by Exo III. The GAA TCA part in the SUV linker is used as a toehold for the SUV releaser. In the cap releaser strand, AGT GCT GA is used as the toehold. Note that Cy5-DNA is a chimeric DNA/RNA oligonucleotide with rArU indicating RNA bases.

| Name | Sequence |
|---|---|
| SUV strand | TAA CAA CCA AAC CAT TTT T /3CholTEG/ |
| GUV strand | /amine/ GGA CAG AGT GAC ATC |
| SUV linker | ATG GTT TGG TTG TTA GAT GTC ACT CTG TCC GAA TCA |
| Hairpin signal containing SUV linker | ATG GTT TGG TTG TTA GAT GTC ACT CTG TCC GAA TCA ACA TCT AAC AAC CAA ACC AT |
| SUV releaser | TGA TTC GGA CAG AGT GAC ATC TAA CAA CCA AAC CAT |
| Cap releaser | TGT CAC TCT GTC CGA ATC AGC ACT |
| Cy5-DNA | /5Cy5/ GGT GGT GGT GGT TGT GGT GGT GGT GGG TCA CTC rArUG TCC GAA TCA GCA CTT TTT TTT TTT |

### Origami design

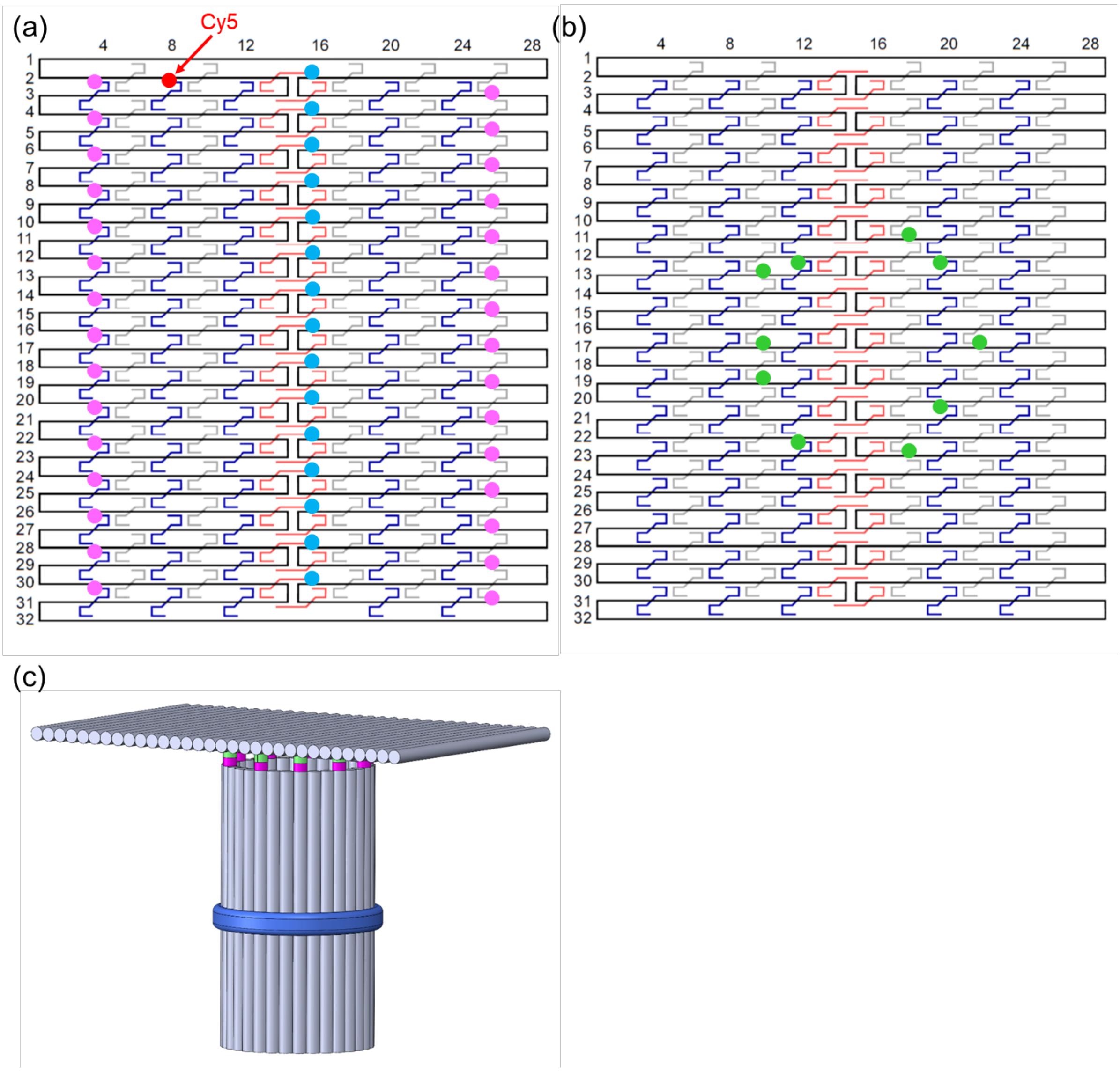


**Figure S1**. The origami designs and folding path diagrams for the tubular pore (a) and rectangular cap (b). (a) The 32-helix origami tile (identical in both (a) and (b)) before cyclization to form the tubular pore (see Table S2 and S3 for staples). A set of staples (termed tubular staples) are used to connect the top and bottom of the tile, forming a tubule (see Table S4). The tubular staples are not shown in the schematic. A Cy5-DNA strand used for fluorescence imaging, shown in Figure 2a-b, is marked in red. A total of 30 staple strands (15 on each side) are added a 16-nt extension to connect the pore with the cap. The 15 staples marked in sky-blue are designed to link with cholesterol-DNA with a 27-nt extension (see Table S3). (b) The rectangular origami cap. Each of 10 staples (indicated by green dots) have a 24-nt extension on 5' end: 16-nt (TTCGGACAGAGTGACA) for binding the pore, and 8-nt (AGTGCTGA) for toehold. (c) Schematic of the tubular pore closed with the rectangular cap. The sky-blue stripe represents the cholesterol moieties. The pink and green denote the staple extensions for capping.

**Table S2**. Blue and gray staple sequences for the origami tile shown in Figure S1. Colored parts correspond to the colored dots in Figure S1. For example, staple [02,08] in Figure S1a has a red dot. In this table, there is a red '/5Cy5/' at the 5' end of the staple's sequence. Similar notations apply to pink (pore-cap connection) and green (cap-pore connection) colored parts. For the staple without a colored dot, colored parts will not show up in the sequence. This means that staple [02,08] in Figure S1b does not have '/5Cy5/' at the 5' end.

| Blue staples | | Gray staples | |
|---|---|---|---|
| Name | Sequence | Name | Sequence |
| [02, 04] | TGTCACTCTGTCCGAAGAACGGTACAGAACAATATTACCGAATACCTA | [03, 05] | GTAATATCCGCCAGAATCCTGAGAGTATAACG |
| [02, 08] | /5Cy5/TATAATCAGAACTCAAACTATCGGATGGATTA | [03, 09] | GAGTAGAAGTGAGGCCACCGAGTAGAGCGGGC |
| [02, 12] | TGTCCATCGATTAGTAATAACATCACACGACC | [03, 17] | AAAATCCCTGAGTGTTGTTCCAGTCGATTTAG |
| [02, 20] | AGAGTCCATTTGATGGTGGTTCCGAGAGGCGG | [03, 21] | AAATCCTGCTATTAAAGAACGTGGAAGCACTA |
| [02, 24] | GTCAAAGGACGCTGGTTTGCCCCATTTTTCTT | [03, 25] | TGTCACTCTGTCCGAAAGCGGTCCGCGAAAAACCGTCTATCAAATCAA |
| [04, 04] | TGTCACTCTGTCCGAACATTTTGAATGCGCGAACTGATAGAACCACCA | [05, 05] | AGTCTTTACGCTCAATCGTCTGAACCTTGCTG |
| [04, 08] | TTTACATTAGACAATATTTTTGAACGGTCAGT | [05, 09] | CGTGGCACGGCAGATTCACCAGTCACTTGCCT |
| [04, 12] | AGTAATAATTCTGACCTGAAAGCGACGCTGAG | [05, 17] | TCCAGTCGCGGCCAACGCGCGGGGAAATCGGC |
| [04, 20] | TTTGCGTATTGCGTTGCGCTCACTGGGTACCG | [05, 21] | CACATTAATTGGGCGCCAGGGTGGGCAGGCGA |
| [04, 24] | TTCACCAGGGGTGCCTAATGAGTGTAGCTGTT | [05, 25] | TGTCACTCTGTCCGAAAAAGCCTGTGAGACGGGCAACAGCTTGCAGCA |
| [06, 04] | TGTCACTCTGTCCGAAGCAGAAGATGAGGAAGGTTATCTATTAGAGCC | [07, 05] | AAAGGAATTAAAACAGAGGTGAGGTGGCTATT |
| [06, 08] | ATTAACACGGTCAGTTGGCAAATCTTTAGAAG | [07, 09] | CAATATCTCGCCTGCAACAGTGCCTAAGAATA |
| [06, 12] | AGCCAGCAAACCTCAAATATCAAACAACTCGT | [07, 17] | AAAACGACACTCTAGAGGATCCCCGCCCGCTT |
| [06, 20] | AGCTCGAAGGGTTTTCCCAGTCACTCTGGTGC | [07, 21] | TAACGCCATTCGTAATCATGGTCAAGCTAACT |
| [06, 24] | TCCTGTGTGTGCTGCAAGGCGATTGCCATTCA | [07, 25] | TGTCACTCTGTCCGAAAGGGGGATGAAATTGTTATCCGCTTAAAGTGT |
| [08, 04] | TGTCACTCTGTCCGAAGTCAATAGCTGATTATCAGATGATATTATACT | [09, 05] | TCATATTCATAATACATTTGAGGAAACAGTTG |
| [08, 08] | TATTAGACCCACCAGAAGGAGCGGCTACCATA | [09, 09] | CAAAGAAATTTACAAACAATTCGACCCTCAAT |

| | | | |
|---|---|---|---|
| [08, 12] | ATTAAATCGAGTAACATTATCATTGAAATAAA | [09, 17] | CGACGACAGCTTTCCGGCACCGCTGACGTTGT |
| [08, 20] | CGGAAACCCGTGCATCTGCCAGTTTAATTCGC | [09, 21] | ATCGTAACAGGCAAAGCGCCATTCAAGTTGGG |
| [08, 24] | GGCTGCGCGGTCACGTTGGTGTAGTCATCAAC | [09, 25] | TGTCACTCTGTCCGAAATGGGATAAACTGTTGGGAAGGGCCTGGCGAA |
| [10, 04] | TGTCACTCTGTCCGAATCTGAATAAAGTTACAAAATCGCGCAAAAGAA | [11, 05] | TGAATACCATGGAAGGGTTAGAACAATTATCA |
| [10, 08] | TCAAAATTAATAACGGATTCGCCTACAAAATT | [11, 09] | GGAGAAACATTTGCACGTAAAACATTGCGGAA |
| [10, 12] | GAAATTGCACAGTAACAGTACCTTAATTACCT | [11, 17] | AGTGCTGATTCGGACAGAGTGACACATTAAATGGAACGCCATCAAAAATGAGGGGA |
| [10, 20] | GTCTGGCCACGTTAATATTTTGTTGGTCATTG | [11, 21] | AATTGTAATTCCTGTAGCCAGCTTATGGGCGC |
| [10, 24] | ATTAAATGGAAGATTGTATAAGCAGAATCGAT | [11, 25] | TGTCACTCTGTCCGAAAAAAACAGTGAGCGAGTAACAACCTGACCGTA |
| [12, 04] | TGTCACTCTGTCCGAAGATGATGAGCGATAGCTTAGATTAAAAATCAT | [13, 05] | AAAACATAAACAAACATCAAGAAAGATTGCTT |
| [12, 08] | AATTACATATTAATTTTCCCTTAGCCTCCGGC | [13, 09] | AGTGCTGATTCGGACAGAGTGACACGCTATTATTAACAATTTCATTTGTTACATCG |
| [12, 12] | AGTGCTGATTCGGACAGAGTGACATTTTTAATAATAACCTTGCTTCTGGTAAATGC | [13, 17] | TGATAAATTCTACAAAGGCTATCAAAAATTCG |
| [12, 20] | AGTGCTGATTCGGACAGAGTGACACCTGAGAGAATATGATATTCAACCTAATACTT | [13, 21] | TCACCATCTCTGGAGCAAACAAGAAATATTTA |
| [12, 24] | GAACGGTATGAGAAAGGCCGGAGACGCAAGGA | [13, 25] | TGTCACTCTGTCCGAACAAAAGGGATCGTAAAACTAGCATAAAGCCCC |
| [14, 04] | TGTCACTCTGTCCGAAAGGTCTGACGTGTGATAAATAAGGTACTAGAA | [15, 05] | ATACCGACGAGACTACCTTTTTAAAATCCTTG |
| [14, 08] | TTAGGTTGCTGACCTAAATTTAATTTATACAA | [15, 09] | TTCATCTTGGTTATATAACTATATTAAATCGT |
| [14, 12] | TGATGCAATTTTTCAAATATATTTCTCAACAG | [15, 17] | AATAAAGCAAACATTATGACCCTGGTTCTAGC |
| [14, 20] | TTGCGGGAGGCAAAGAATTAGCAAGTAGATTT | [15, 21] | ACAGGCAAGAAGCCTTTATTTCAACAGTCAAA |
| [14, 24] | TAAAAATTTAGCATTAACATCCAACAAATGGT | [15, 25] | TGTCACTCTGTCCGAAAATAGTAGTTTAGAACCCTCATATTAAAGATT |
| [16, 04] | TGTCACTCTGTCCGAAAAAGCCTGAAAGTAATTCTGTCCAGCAGAACG | [17, 05] | CAAAAGGTTTTAGTATCATATGCGGGTTTGAA |
| [16, 08] | ATTCTTACTAATAAGAGAATATAAGTCCTGAA | [17, 09] | AGTGCTGATTCGGACAGAGTGACACGAGCCAGCAGTATAAAGCCAACGTAGTTAAT |
| [16, 12] | TAGGGCTTATGTAATTTAGGCAGAATTTACGA | [17, 17] | TACGGTGTCCAATTCTGCGAACGAAATTAAGC |

| | | | |
|---|---|---|---|
| [16, 20] | AGTTTGACCATGTTTTAAATATGCTAATTCGA | [17, 21] | AGTGCTGATTCGGACAGAGTGACATAGCTCAACATTAGATACATTTCGTAAATCAT |
| [16, 24] | CAATAACCGCTTAATTGCTGAATAGCAAACTC | [17, 25] | TGTCACTCTGTCCGAAGGCTTAGATGTTTAGCTATATTTTATTCTACT |
| [18, 04] | TGTCACTCTGTCCGAACGCCTGTTTTCATCGTAGGAATCAAGAAGGCT | [19, 05] | TTTTTATTTATCAACAATAGATAAAGTACCGA |
| [18, 08] | CAAGAAAACACTCATCGAGAACAAGCGTTTTA | [19, 09] | AGTGCTGATTCGGACAGAGTGACAAAGTACCGATAATATCCCATCCTAGGCATTTT |
| [18, 12] | GCATGTAGCATTCCAAGAACGGGTTTTTGAAG | [19, 17] | GCGGATTGTTCAAATATCGCGTTTAACTAAAG |
| [18, 20] | GCTTCAAACTGACTATTATAGTCAGCCAGAGG | [19, 21] | TCTTTACCGCGAACCAGACCGGAATAATGCTG |
| [18, 24] | CAACAGGTAATGACCATAAATCAAGGATAGCG | [19, 25] | TGTCACTCTGTCCGAAAAAACGAGCAGGATTAGAGAGTACTTGCGGAT |
| [20, 04] | TGTCACTCTGTCCGAATATCCGGTAATAAACAGCCATATTTGTTTAAC | [21, 05] | AGTTACAAATTCTAAGAACGCGAGGCAAGCCG |
| [20, 08] | GCGAACCTCGTCTTTCCAGAGCCTTACAGAGA | [21, 09] | CTAACGAGCCCGACTTGCGGGAGGATTAAACC |
| [20, 12] | CCTTAAATATTTTATCCTGAATCTCATTAGAC | [21, 17] | TTTACCAGTTGCAAAAGAAGTTTTGAAGCAAA |
| [20, 20] | AGTGCTGATTCGGACAGAGTGACAGGGTAATAAGAGCAACACTATCATCGTTGGGA | [21, 21] | GCATAGTAGTAAAATGTTTAGACTAAATCAGG |
| [20, 24] | TCCAATACCATAACGCCAAAAGGAAACTAACG | [21, 25] | TGTCACTCTGTCCGAATGCAGATATGCGGAATCGTCATAACAGTTCAG |
| [22, 04] | TGTCACTCTGTCCGAAGTCAAAAAAAACAATGAAATAGCAAGTAAGCA | [23, 05] | AGAGCAAGTGAAAATAGCAGCCTTAATTTGCC |
| [22, 08] | GAATAACAAGAATTGAGTTAAGCCGGAAACCG | [23, 09] | AACCCACATAAAAACAGGGAAGCGTACCAACG |
| [22, 12] | AGTGCTGATTCGGACAGAGTGACAGGGAGAATATTGAGCGCTAATATCCCCAAAAG | [23, 17] | AGTGCTGATTCGGACAGAGTGACAATCATTGTATTATACCAGTCAGGAAACCCTCG |
| [22, 20] | AGAAAAATTGAGATGGTTTAATTTGGCGCATA | [23, 21] | ATTGGGCTCTACGTTAATAAAACGATTACGAG |
| [22, 24] | GAACAACAGAGAAACACCAGAACGAATCTTGA | [23, 25] | TGTCACTCTGTCCGAAGCCCTGACTTATTACAGGTAGAAATCAACTAA |
| [24, 04] | TGTCACTCTGTCCGAAGATAGCCGTAAGTTTATTTTGTCAGCCAAAGA | [25, 05] | CCACGGAAAACAAAGTTACCAGAACAATAATA |
| [24, 08] | AGGAAACGACATATAAAAGAAACGGAGGGAGG | [25, 09] | AGGTGGCACAATAATAACGGAATAAGAGAGAT |
| [24, 12] | AACTGGCACAAACGTAGAAAATACTTCATTAA | [25, 17] | TAAGGGAAAACGGTGTACAGACCACAACTTTA |
| [24, 20] | GGCTGGCTGAACGAGGCGCAGACGCGAAAGAG | [25, 21] | CTTAGCCGGACCTTCATCAAGAGTAGTAGTAA |

| | | | |
|---|---|---|---|
| [24, 24] | CAAGAACCAAATCCGCGACCTGCTATCTTTGA | [25, 25] | TGTCACTCTGTCCGAATTGTGTCGGGATATTCATTACCCATAAGGCTT |
| [26, 04] | TGTCACTCTGTCCGAACAAAAGGGAACCATCGATAGCAGCCTTTAGCG | [27, 05] | ACCAATGACGACATTCAACCGATTCAAAGACA |
| [26, 08] | GAAGGTAAACCATTAGCAAGGCCGGCATTTTC | [27, 09] | GCACCATTATATTGACGGAAATTAATACATAA |
| [26, 12] | AGGTGAATTTAGAGCCAGCAAAATTGCCATCT | [27, 17] | AGTTTCCAAGGCACCAACCTAAAAGTCAATCA |
| [26, 20] | GCAAAAGAGGACTAAAGACTTTTTTGACAACA | [27, 21] | GGCTTTGAATACACTAAAACACTCCCATGTTA |
| [26, 24] | CCCCCAGCAACGAGGGTAGCAACGTATTCGGT | [27, 25] | TGTCACTCTGTCCGAAAGCATCGGGATTATACCAAGCGCGCTGATAAA |
| [28, 04] | TGTCACTCTGTCCGAATCAGACTGCCACCAGAACCACCACGGCAGGTC | [29, 05] | CAGAGCCGTAGCGCGTTTTCATCGGAAACGTC |
| [28, 08] | GGTCATAGCGCCACCCTCAGAGCCAACAAATA | [29, 09] | CTCAGAACCCCCCTTATTAGCGTTCACCAGTA |
| [28, 12] | TTTCATAAACCGCCTCCCTCAGAGAAGCGCAG | [29, 17] | GCTTGCTTATAGTTGCGCCGACAACATGAGGA |
| [28, 20] | ACCATCGCGCCTTTAATTGTATCGTTAGTAAA | [29, 21] | CAAAAGGACCACGCATAACCGATAGCTACAGA |
| [28, 24] | CGCTGAGGTGAAAATCTCCAAAAATAAACAAC | [29, 25] | TGTCACTCTGTCCGAATTTCACGTCTTGCAGGGAGTTAAACGAAAGAC |
| [30, 04] | TGTCACTCTGTCCGAAAGACGATTCGTATAAACAGTTAATAAACATGA | [31, 05] | ACAGTGCCGGCCTTGATATTCACAACCACCCT |
| [30, 08] | AATCCTCATTAACGGGGTCAGTGCCAAGAGAA | [31, 09] | AATAAGTTTTAAAGCCAGAATGGACCGCCACC |
| [30, 12] | TCTCTGAATTGATGATACAGGAGTTCAGTACC | [31, 17] | ACAGACAGGTCGTCTTTCCAGACGGTTTATCA |
| [30, 20] | TGAATTTTAAACTACAACGCCTGTCACCGTAC | [31, 21] | ACCAGTACCTGTATGGGATTTTGCAAAGGCTC |
| [30, 24] | TTTCAACATGTACCGTAACACTGATCAGAACC | [31, 25] | TGTCACTCTGTCCGAAGGAACCCAGTTTCAGCGGAGTGAGAATAATTT |

**Table S3**. Red staple sequences for the origami tile shown in Figure S1. Sky-blue parts correspond to the sky-blue dots in Figure S1. For example, staple [02,16] in Figure S1a has a sky-blue dot. In this table, there is a sky-blue extension at the 3' end of the staple's sequence. All the sky-blue extensions are designed to bind with a cholesterol-DNA, whose sequence is TGGACGGCCGTCAACTGCGGCGTGTAA/3CholTEG/. For the staple without a sky-blue dot, colored parts will not show up in the sequence. This means, for example, that staple [02,16] in Figure S1b does not have the blue extension at the 3' end.

| Red staples | | | |
|---|---|---|---|
| Name | Sequence | Name | Sequence |

| | | | |
|---|---|---|---|
| [02, 16] | GATAGGGTTTATAAATCAAAAGAAGTAGCAATTTACACGCCGCAGTTGACGGCCGTCCA | [17, 13] | ACGCCAACAATTGAGAATCGCCATATATAACA |
| [03, 13] | ACTTCTTTACGCAAATTAACCGTTTAGCCCGA | [18, 16] | CGAAAGACCATCAAAAAGATTAAGGGCTGTCTTTACACGCCGCAGTTGACGGCCGTCCA |
| [04, 16] | TAATGAATGGAAACCTGTCGTGCCAACAGAGATTACACGCCGCAGTTGACGGCCGTCCA | [19, 13] | TTCCTTATAAACCAATCAATAATCAGGAAGCC |
| [05, 13] | TAGAACCCAAGGGACATTCTGGCCAGCTGCAT | [20, 16] | AGAGGCTTACGACGATAAAAACCATTTGCACCTTACACGCCGCAGTTGACGGCCGTCCA |
| [06, 16] | GCAGGTCGGGCCAGTGCCAAGCTTAAGCATCATTACACGCCGCAGTTGACGGCCGTCCA | [21, 13] | CAGCTACACAAGATTAGTTGCTATAAATAGCG |
| [07, 13] | CCTTGCTGGCAAATGAAAAATCTAGCATGCCT | [22, 16] | ACTGGCTCGAATTACCTTATGCGAACAAAGTCTTACACGCCGCAGTTGACGGCCGTCCA |
| [08, 16] | TCCAGCCAGTATCGGCCTCAGGAATTAATTTTTTACACGCCGCAGTTGACGGCCGTCCA | [23, 13] | AGAGGGTATAACTGAACACCCTGATTTTAAGA |
| [09, 13] | AAAAGTTTCTTTGCCCGAACGTTAGATCGCAC | [24, 16] | GACAGATGCCGAACTGACCAACTTTTACGCAGTTACACGCCGCAGTTGACGGCCGTCCA |
| [10, 16] | AACCAATATTTTGTTAAATCAGCTAACGTCAGTTACACGCCGCAGTTGACGGCCGTCCA | [25, 13] | TATGTTAGTGATTAAGACTCCTTATGAAAGAG |
| [11, 13] | ATGAATATGTAGATTTTCAGGTTTCATTTTTT | [26, 16] | CACTACGATTAAACGGGTAAAATATTGAGCCATTACACGCCGCAGTTGACGGCCGTCCA |
| [12, 16] | TTGAGAGATAATGCCGGAGAGGGTTCAATATATTACACGCCGCAGTTGACGGCCGTCCA | [27, 13] | TTTGGGAATATCACCGTCACCGACCGTAATGC |
| [13, 13] | TGTGAGTGGGAAACAGTACATAAAAGCTATTT | [28, 16] | TGATACCGTCGAGGTGAATTTCTTCAGAGCCATTACACGCCGCAGTTGACGGCCGTCCA |
| [14, 16] | TGTACCAACTCAGAGCATAAAGCTAAGAACGCTTACACGCCGCAGTTGACGGCCGTCCA | [29, 13] | CCACCGGATCAAAATCACCGGAACAAACAGCT |
| [15, 13] | GAGAAAACATCCAATCGCAAGACAAAATCGGT | [30, 16] | AAAGTTTTCCCTCATAGTTAGCGTGCGTCATATTACACGCCGCAGTTGACGGCCGTCCA |
| [16, 16] | GTTGATTCCTGGAAGTTTCATTCCATTTAACATTACACGCCGCAGTTGACGGCCGTCCA | [31, 13] | CATGGCTTTTTACCGTTCCAGTAAAACGATCT |

**Table S4**. Sequence of tubular staples. These staples are designed to bind the upper and lower boundaries of the origami tile together, thus forming a tubular pore.

| Tubular staples | | | |
|---|---|---|---|
| Name | Sequence | Name | Sequence |
| [1,8] | AACAGGAGGGAACCTATTATTCTGGCCCCCTG | [32,25] | AAGTATTATCGTTAGAATCAGAGCTTAGACAG |
| [1,40] | TGCTTTCCAGAGGCTGAGACTCCTCTTGAGTA | [32,57] | GGATTAGGGCTGGCAAACGAGCACAGTGTTTT |
| [1,72] | GCTAGGGCATTAGCGGGGTTTTGCGTACTGGT | [32,89] | AGGCGGATGGGAAGAAAGCGAAAGAAAGAGTC |

| | | | |
|---|---|---|---|
| [1,104] | GAAAGGAAAAGTGCCGTCGAGAGGGTTGATAT | [32,121] | AAGTATAGGGGGAAAGCCGGCGAACGTGGCGA |
| [1,136] | AGCTTGACCCCGGAATAGGTGTATAGCATTCC | [32,153] | TCAGGAGGCCCTAAAGGGAGCCCCTTGGAACA |
| [1,168] | AATCGGAATTTAGTACCGCCACCCGTTTCGTC | [32,185] | GCCACCCTGGGTCGAGGTGCCGTAACTCCAAC |
| [1,200] | GTTTTTTGCAGAACCGCCACCCTCGCCCAATA | [32,217] | CACCCTCATACGTGAACCATCACCCAGGGCGA |

**Lipid information**

All the lipids used are purchased from Avanti Polar Inc.

Lipids in SUV

1. 16:0 PC or DPPC: 1,2-dipalmitoyl-sn-glycero-3-phosphocholine
2. 16:0 Liss Rhod PE: 1,2-dipalmitoyl-sn-glycero-3-phosphoethanolamine-N-(lissamine rhodamine B sulfonyl) (ammonium salt)

Lipids in GUV

1. 14:0 PC or DMPC: 1,2-dimyristoyl-sn-glycero-3-phosphocholine
2. 18:1 Biotinyl PE: 1,2-dioleoyl-sn-glycero-3-phosphoethanolamine-N-(biotinyl) (sodium salt)
3. DSPE-PEG(2000)-DBCO: 1,2-distearoyl-sn-glycero-3-phosphoethanolamine-N-[dibenzocyclooctyl(PEG)-2000]

### 2. Sample Preparation Methods

#### Conjugation of DNA to lipid molecules

DNA-lipid conjugates are used as components of DNA-decorated GUVs. The method for conjugation is as described in a previous report.[1] The method is a two-step process. The first step is a modification of DNA with azide group. The amine modified DNA strands in Table S1 were synthesized with azide-N-hydroxysuccinimide or azide-NHS in dimethylformamide or DMF, and triethylamine (TEA) was added as catalyst. The mixture was incubated at room temperature in shades for two hours to achieve complete reaction. Then, we added 200 µL ethanol and 10 µL 4 mM NaCl into the solution and froze it at -20 °C for 30 minutes to precipitate DNA. The solution was then centrifuged at 20,000 g for 30 minutes. The precipitate was re-dispersed in 200 µL ethanol and centrifuged for several times to remove excess azide-NHS molecules. After centrifugation, the precipitate was dried in vacuum and re-suspended in 1x phosphate buffered saline (PBS). The concentration of synthesized DNA-azide was determined by the absorption of the sample at 260 nm using a Perkin-Elmer Lambda 950 UV/visible/NIR spectrophotometer.

The second step is a conjugation of azide-DNA with DSPE-PEG(2000)-DBCO. DBCO can react with azide by click chemistry. DNA-azide from the last step was mixed with DSPE-PEG(2000)-DBCO at a molar ratio of 1:5. The mixture was then purified with centrifugation and ready for use.

#### Polystyrene particles with Exo III

Approximately 5 µL 10 µM Exo III enzyme (New England Biolabs) was mixed with 2% polystyrene particle solution (Thermo Fisher Scientific) diluted in 70 µL 2-(N-morpholino) ethanesulfonic acid (MES) buffer (pH 6.0). Then, 5 µL 100 µM 1-ethyl-3-(3-dimethylaminopropyl)carbodiimide (EDC) was added to the mixture as catalyst. In order to keep the activity of Exo III molecules, the mixture was kept at 4 °C for 3 hours. After the reaction was completed, the mixture was purified with the centrifugation method to remove the excess enzymes. The extinction coefficient of polystyrene particle is approximately $2x10^9$ $M^{-1}cm^{-1}$ at 660nm. It is measured that the polystyrene concentration is 0.19nM. Exo III is estimated around 1nM. Therefore, an average of 5 Exo III enzyme attached per polystyrene particle.

The enzymatic activity of Exo III after conjugation with polystyrene particles was examined by measuring the change of the fluorescence peak intensity from a DNA duplex modified with fluorescent dyes. One strand was conjugated with FAM dye (5' – ATC GGT CAG GCT T/iFluorT/T TTTTTT T – 3') and the other has a quencher on its 5' end (5' – /5IABkFQ/ AAG CCT GAC CGA T – 3'). When the two strands are hybridized, the quencher can absorb the fluorescence from the FAM dye, thus no fluorescence will be observed. If the duplex is digested partly by the enzyme, strong fluorescence will emerge. To test the Exo III activity, 1 µM of each DNA strand was dissolved in 100 µL 1x NEB Buffer (B7001S, New England Biolabs) first and measured fluorescence. Then, the measurement was paused, 1 µL conjugated Exo III particle solution was added into the solution, and the measurement was resumed. From the change of FAM fluorescence, the activity of Exo III was evaluated.

**Assembly of DNA origami structures**

As described in Figure S1, two DNA origami structures were used in this study. A tubular origami was designed as a transmembrane pore, while the rectangular tile was used for capping the pore. The design of the DNA tubule is identical with that of the origami rectangle except for the use of tubular staples.[2]

The origami tubules were prepared by mixing 10 nM scaffold strands, 4× DNA staples (Tables S2 and S3), 14× tubular staples (Table S4), and 160× cholesterol modified DNA in 1× TAEM buffer (an aqueous solution of 40 mM trisaminomethane, 1 mM ethylenediaminetetraacetic acid (EDTA) disodium salt, 20 mM acetic acid, and 10 mM $MgCl_2$ at pH ~8). Note that cholesterol DNA is used for the insertion into lipid membranes. The mixture was then annealed in a Bio-Rad S1000 thermal cycler from 75 °C to 4 °C at -1 °C per minute.

The origami tiles were synthesized by mixing 10 nM scaffold strands with 4× DNA staples (Tables S2 and S3) in 1× TAEM buffer. The annealing of the mixture went from 75 °C to 4 °C at -1 °C per minute in the thermal cycler.

The origami tubules and tiles were purified 3 times by using the centrifugal filter (100 kDa) from Amicon. Then, the purified tubules and tiles were mixed at 1:2 molar ratio for assembly of capped origami pores. After that, the mixture was annealed from 55 °C to 4 °C at -1 °C per minute in the thermal cycler.

**Synthesis of giant vesicles**

DNA-lipid molecules synthesized from the previous step was mixed with DMPC at a molar ratio of 1:1000 in a glass vial.[3-5] The solution was then dried in vacuum for 30 minutes to evaporate all solvent and resuspended with 600 μL liquid paraffin. The new solution was sonicated at 50 °C for 3 hours. Lipids will disperse uniformly in the sonicated solution. Then, 10 μL 10 nM DNA origami tubules, 5 μL Exo III-particle, both in 1× TAEM were mixed and additional TAEM buffer added to adjust the volume of the mixture to 20 μL. This mixture was added into the liquid paraffin containing lipids and vortexed for 25 seconds to form aqueous droplets. After vortex, the vesicle solution became blurred. Then, 600 μL of this vesicle solution was poured onto 300 μL TAEM buffer and centrifuged for 15 minutes at 8,000 g. The giant vesicles including DNA strands, origami tubules, and Exo III-particles were in the precipitates. Both aqueous phase and oil phase supernatants were discarded, and the precipitate was dissolved in TAEM buffer for future use. From the molar ratio of components, we estimate that there are about 3,000,000 DNA-lipid conjugates integrated on the surface of each giant vesicle. For the giant vesicles without membrane pores, the tubular origami was not included in the synthesis.

**Small vesicles with DNA strands**

Small vesicles with DNA strands were prepared using a dehydration-rehydration method.[6] Cholesterol modified DNA (*i.e.*, SUV strands in Table S1) and rhodamine B functionalized lipid (*i.e.*, 16:0 Liss Rhod PE) were mixed with DPPC at a molar ratio of 1:1:1000 in a glass vial.

The solution was dried in vacuum for 20 minutes to let lipids form a dry thin film on the bottom of the vial. Then, 1 mL TAEM buffer was added to the glass vial and the solution was placed on a pre-heat hot plate at ~90 °C. The solution was stirred with a stirring bar at 500 rpm for 1 hour while the temperature was kept all the time. After stirring, small vesicle solution was purified by centrifugation with a 30 kDa molecular weight cut off spin column at 5,000 g for 5 minutes. This process was repeated 6 times to remove unbound lipid and DNA molecules. From the molar ratio of components, we estimate roughly 50 oligonucleotides per small vesicle.

### 3. Experimental Setup

#### Flow channel assembly

The experiments were conducted with a multi-channel flow cell assembled with a piece of glass coverslip (Schott) and a quartz slide sealed with medical grade acrylic adhesive sheets. The estimate channel volume is approximately 20 μL. Inlet and outlet ports (LabSmith) were glued to the glass slides using epoxy. In the experiments, Tygon microbore tubing was used to connect sample tubes and the flow channel. The assembled microfluidic channel was placed under an inverted fluorescence microscope for optical imaging.

#### Surface passivation

The piranha washed glass coverslips were passivated with a one-step method to coat BSA-biotin on the coverslip and prevent nonspecific interactions between the vesicles and the glass surface. Before the experiments, approximately 30 μL 5 μM BSA-biotin in TAEM and tween-20 solution was flown into the channel and incubated for 1 hour. Tween-20 was used to fix defects on the coverslip surface, and BSA-biotin was attached to the surface in order for biotin-streptavidin conjugation that immobilizes giant vesicles on the substrate. After the passivation processes, ~30 μL 1 μM streptavidin in TAEM buffer was added to the fluid channel. Without adding streptavidin, the GUVs with biotin moieties will not be attached on the surface.

#### Imaging system

A custom-built inverted fluorescence microscope (Zeiss Axio Observer D1) was used for imaging. Three diode lasers at 405, 561, and 658 nm (Laserglow) were used as light source. An oil-immersion 63× objective lens from Zeiss was used, and the collected emission light from the sample was imaged with an Andor iXon3 electron multiplying charge coupled device (EMCCD) camera.

### 4. Characterization

#### AFM images of tubular and rectangular DNA origami

We conjugated origami tiles with DNA tubules with a molar ratio of 2:1 to ensure as many origami pores capped as possible. We performed AFM imaging to examine whether the origami tubules are closed with flat origami caps as designed in Figure S1. For deposition of samples for AFM imaging, the target sample was diluted to 1.0 nM with TAEM buffer. Then, an aliquot of 10 μL diluted sample was pipetted onto mica surface for 5-minute incubation at room temperature. After that, the mica was blown dry with compressed air, rinsed with 80 μL deionized (DI) water for about 3 seconds, and then blown dry again with compressed air.

AFM imaging was performed in air using the Peak-Force tapping mode with a Bruker Dimension Icon AFM and SCANASYST-AIR probes. The images show that more than 90% of the examined tubules are connected with flat rectangular tiles (Figure S2). The connection is also confirmed by the height profiles. After the cap releasers were incubated for 1 hour, nearly 90% of the tubules are detached from the flat origami, confirming that the cap releasers are effective in removing the caps and opening the origami pores.

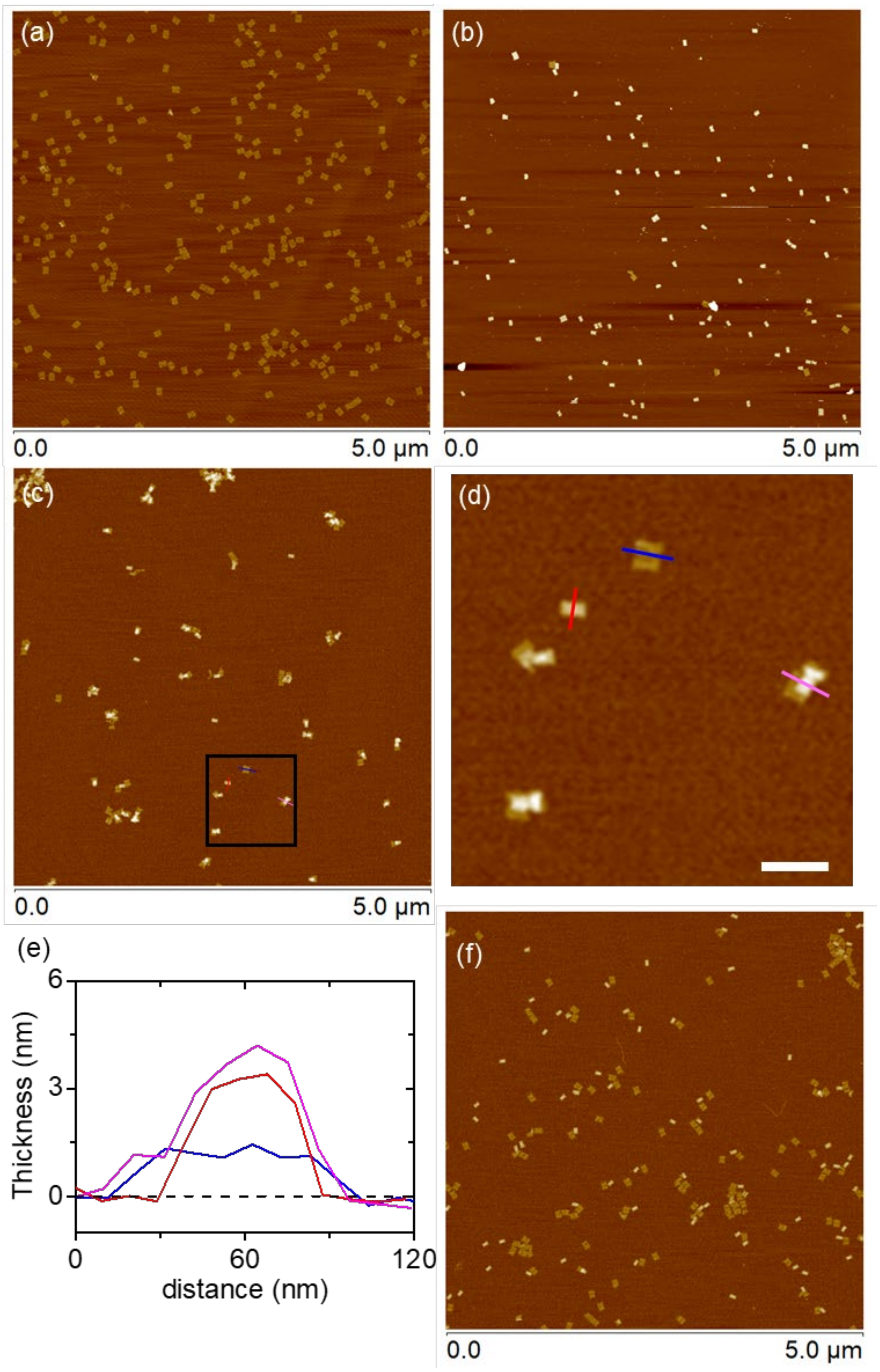


**Figure S2**. AFM images of DNA origami. (a) Near 100% flat origami rectangles, which measure approximately 60 nm x 100 nm in dimension. (b) The majority of DNA origami measures approximately 60 nm x 50 nm with a thickness of ~4 nm, which is roughly half the size and twice the thickness of the origami rectangle. The results indicate the identity of the tubular structure of the origami in solution, which should have a diameter of ~32 nm and a length of ~60 nm. Note that the tubular structures collapsed into a rectangular shape to maximize their contact with the mica substrate. (c) Origami structures after connecting tubules with tiles at a ratio of 1: 2. Most tubules are connected to flat tiles, while the linked origami structures in 3D collapsed before or during the AFM imaging. (d) Zoom-in of the black square area in (c). The scale bar is 20 nm. Colored lines (blue, red, and pink) denote the sites where heights are measured. (e) Corresponding height profiles of the objects in (d). Cross-sections of a flat tile (blue), an isolated tubule (red), and a tubule connected with a flat tile (pink). Dashed line indicates the mica substrate. The profiles confirm the connection between the tubule and flat tile. (f) Origami tiles and tubules after mixing with cap releaser strands. The majority (~90%) of the tubules are separate from the tiles, suggesting that cap releasers are effective in removing the flat caps from the tubules.

**Additional kinetic measurement**

To measure the kinetics of molecular transport via origami channel, we used GFP and Cy5-DNA in the outflow experiment (Figure 3 and Figure S3). GFP and Cy5-DNA were initially encapsulated inside the giant vesicles at a concentration of ~2 μM. The origami channels were initially closed with the flat origami tiles. The fluorescence intensity did not change without adding cap releaser strands to open the origami pores, indicating no significant leak of fluorescent molecules from the GUV. After ~25 mins of the observation, we added the cap releasers as indicated by black arrows in Figure S3b and S3f. Shortly after that (roughly 5 mins), the fluorescence intensity inside the GUV started to drop. The intensity continued to decrease over time. However, the intensity inside the GUV was still higher than the background after 90 mins. In the control experiments (Figure S3c, S3d, S3g, S3h), the giant vesicles contained the fluorescent molecules, but did not include origami pores. As expected, no significant changes in the fluorescence intensity were observed, indicating that there will be no molecular diffusion in and out of the vesicle without origami channels.

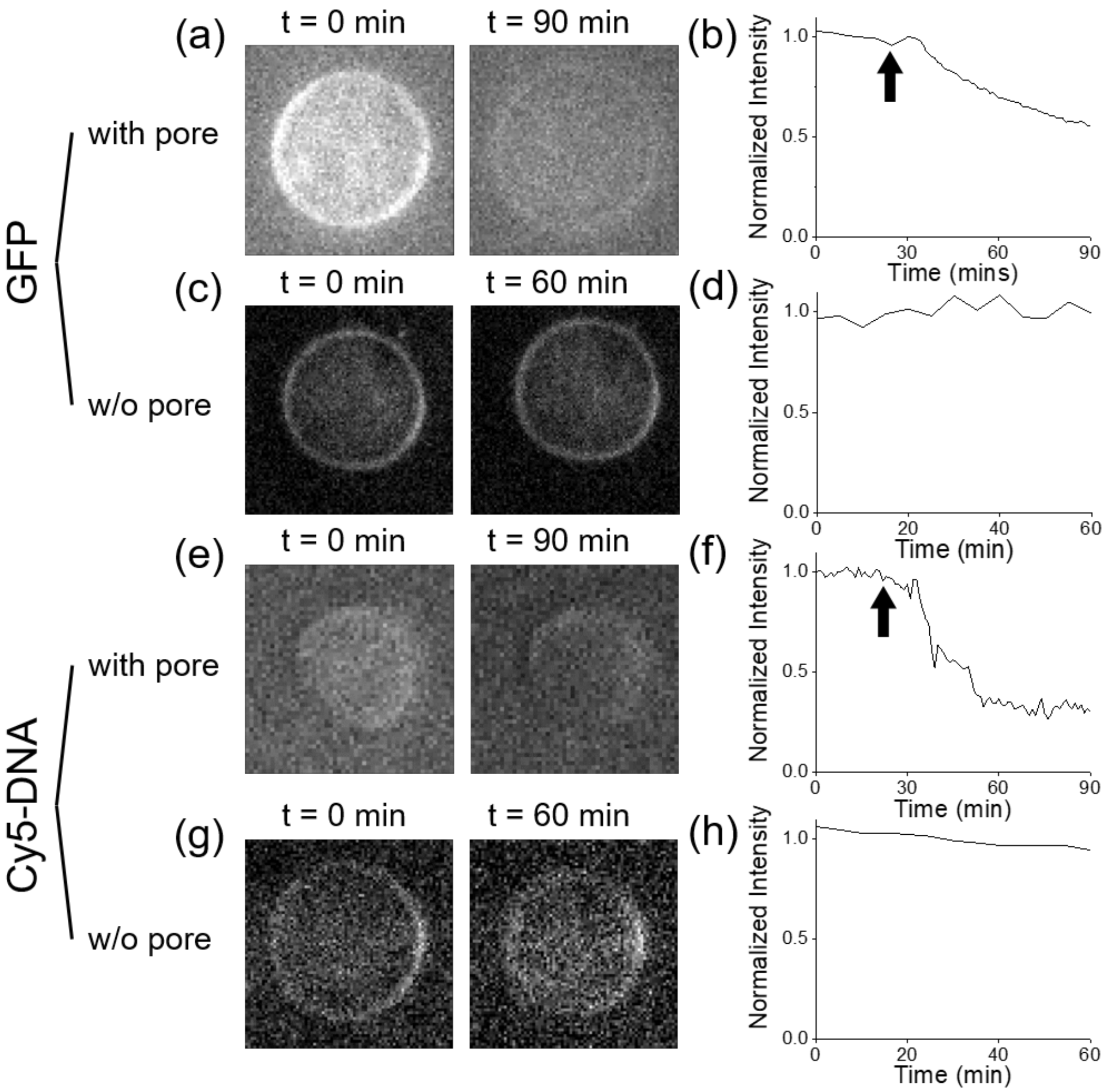


**Figure S3**. Kinetic measurements of molecular transport through DNA origami pores. (a) Fluorescence images of GFP molecules initially encapsulated inside a GUV at the beginning ($t$ = 0 min) and end ($t$ = 90 min) of the experiment. (b) Normalized fluorescence intensity of GFP as a function of time. After 25 mins, the cap releaser strands were introduced to the imaging chamber to remove the origami caps and open the channels, as indicated by the black arrow. Shortly after that, the fluorescent molecules diffused out of the GUV, and the fluorescence intensity inside GUV dropped. (c)-(d) Control experiment with the GUV containing GFP, but without origami pores. The fluorescence intensity does not change significantly, indicating that no molecular transport in and out of the vesicle without pores. (e)-(f) Fluorescence measurement of Cy5-DNA encapsulated inside a giant vesicle over time. Fluorescence images at the beginning and end of the experiment ($t$ = 0 and 90 min, respectively). Shortly after adding cap releaser strands (black arrow), the fluorescence intensity inside the GUV dropped. (g)-(h) Control experiment with the GUV encapsulating Cy5-DNA without origami channels. The fluorescence intensity remains constant over time without origami pores.

**Exo III activity inside a giant vesicle**

In our experiment, Exo III enzymes are used to transduce external DNA signals to another form of signals inside a giant vesicle. For example, the hairpin strand was partly digested by the enzyme, which exposes the SUV linker domain. The enzymes are functionalized on the surface of polystyrene particles with a diameter of ~200 nm. With the particle, the enzymes will not pass through the origami channels that have a diameter of about 32 nm. To prove that Exo III is functional inside the giant vesicle, we examined Exo III activity with both enzymes in free solution and particle-enzymes inside a GUV using a fluorophore-quencher pair. We used a FAM modified strand (termed FAM-DNA) and its complementary strand with a quencher group (quencher-cDNA), as discussed above. When hybridized, the quencher will absorb the emitted fluorescence from the FAM-DNA. For testing the free Exo III enzymes, we first prepared a 1 μM FAM-DNA solution which has a high fluorescence intensity. Then, we added the solution of cDNA-quencher at a ratio of 1:1, and the fluorescence intensity immediately dropped. Next, we added 1 nM Exo III that will digest the cDNA-quencher. The fluorescence intensity increased (Figure S4a), indicating that Exo III cleaved off cDNA and FAM-DNA is thus released.

We also examined the activity of the particle-modified Exo III enzymes and repeat the previous experiment. Again, we observed a sharp increase of the fluorescence intensity after adding enzyme-particles to the FAM-DNA/cDNA-quencher solution, as shown in Figure S4b. Next, we encapsulated the particle-Exo III into the giant vesicles with open origami pores. This time, the fluorescence intensity increase was small, yet distinct upon addition of GUVs containing enzyme-particles (Figure S4c and S4d). We note that the degree of fluorescence recovery upon addition of Exo III varies significantly. For example, the fluorescence recovery with particle-modified enzymes is much less than that with free enzymes, and it is even smaller with particle-enzymes inside giant vesicles. This may be attributed to the fact that the amount of enzymes was reduced significantly during the processes of conjugation, assembly, encapsulation, and purification. Further, the diffusion of oligonucleotides into and out of vesicles through origami channels may present another barrier. Nonetheless, our experiment clearly demonstrates that DNA signals can received, transduced, and transmitted by synthetic cells, which is used to program reversible aggregation behaviors.

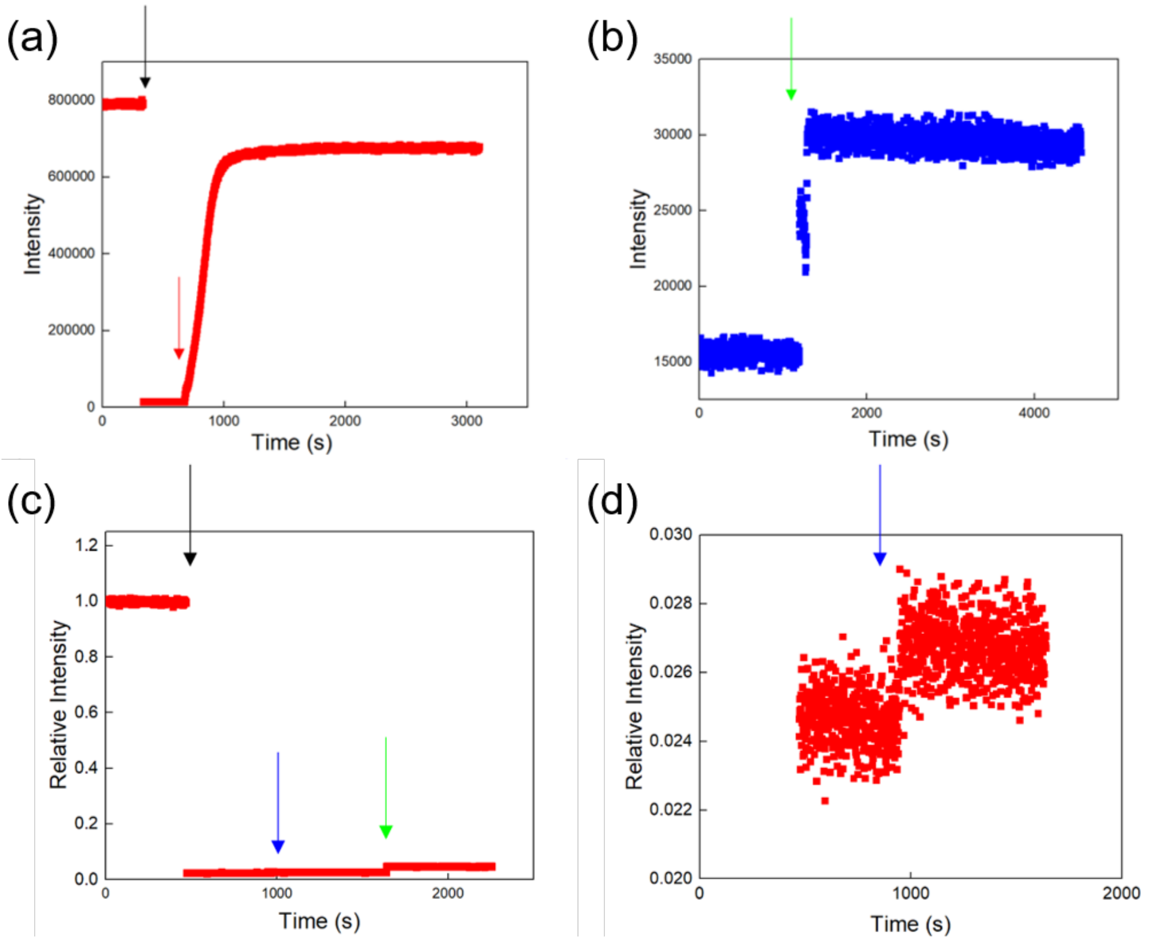


**Figure S4**. Exo III activity measured with a FAM-quencher pair in solution. (a) Fluorescence measurement with free Exo III enzymes. Initially, fluorescence is measured from 1 μM of FAM-DNA in solution. Upon addition of quencher-cDNA at 1 μM (black arrow), the fluorescence intensity immediately drops. After a few minutes, we added 1 nM Exo III enzymes into the solution (red arrow), which results in the increase of fluorescence intensity. (b) Fluorescence measurement in a similar experiment with particle-Exo III rather than free enzymes. The addition of polystyrene particle modified Exo III enzymes into the solution of FAM-DNA and cDNA-quencher (1 μM each; green arrow) leads to the increase of fluorescence intensity. (c)-(d) Fluorescence measurement with enzyme-particles encapsulated in giant vesicles with open origami pores. Black Arrow indicate the addition of cDNA-quencher into the solution of FAM-DNA (1 μM each). Then, GUVs containing enzyme-particles are added to the solution (blue arrow). A small, yet distinct increase of the fluorescence intensity is observed. After a few minutes, particle-Exo III (without vesicles) are added directly to the solution (green arrow), which results in a further increase of the fluorescence intensity.

As Exo III can digest DNA in a duplex from the 3’ end, it is important to measure how it will interact with origami channels. We incubated the particle-modified Exo III with DNA origami tiles for 15 mins, and then performed the AFM imaging. Note that 15-min incubation was used because it takes about 15 mins to encapsulate particle-Exo III in the giant vesicles with DNA origami channels. Figure S5 shows that most of the origami rectangles maintain their original shape. The results strongly suggest that the Exo III will not damage origami structures significantly under the experimental conditions.

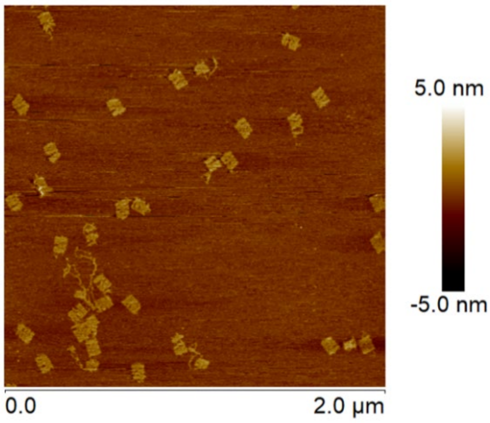


**Figure S5**. AFM image of DNA origami tiles after incubation with Exo III. Approximately 10 nM DNA origami tiles were mixed with 0.1 nM Exo III enzymes in 10 μL TAEM buffer for 15 mins. The AFM image shows that most of the rectangular origami remains its original shape, indicating that this concentration of Exo III will do little damage to the origami.

**Immobilized giant vesicle shape change over time**

During the microscope observation, we find that several the vesicles that change the shapes and sizes over time. Our observation suggests that the immobilized vesicles gradually flatten over time as additional binding occurs between the vesicles and the surface. Further, we also noted that the flow in the imaging chamber may cause some change in the vesicle shape as shown below in Figure S8.

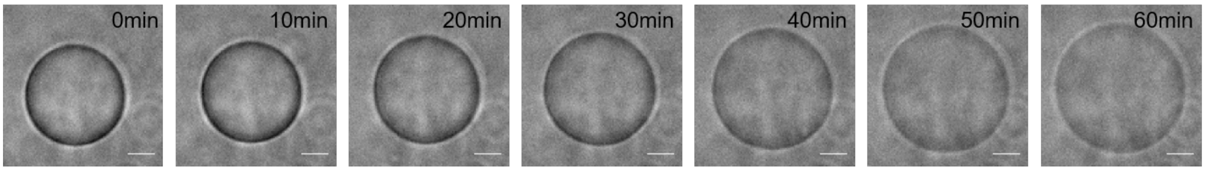


**Figure S6**. Shape change of a GUV during a one-hour period. Scale bar = 5 μm.

**Reversible aggregation behavior using DNA signals**

The reversible aggregation experiment using the SUV linkers was more efficient than using the hairpin strands, because it skips the process of (i) pore opening, (ii) hairpin penetration into the vesicle, (iii) cleavage by Exo III, and (iv) outflow of the transduced oligonucleotides (*i.e.*, exposed SUV linkers) from the GUV. Therefore, we observed more drastic change in fluorescence intensity in a less amount of time. In the aggregation experiment using SUV linkers, approximately 30 mins were given for the SUVs to aggregate onto the giant vesicle, and about 20 mins for the small vesicles to dissociate from the GUV. With the hairpin strands, however, an hour was given for each step during the course of experiment. Figures S7 and S8 present additional reversible aggregation experiments (similar to Figure 5). To confirm that the hairpin strand itself cannot trigger the aggregation, we performed a control experiment that did not include the first step of adding cap releaser strands. Without cap releasers, the origami channels on the giant vesicles remain closed with the rectangular origami caps. As a result, SUVs and hairpins cannot initiate the aggregation of small vesicles on the giant vesicle, as shown in Figure S9.

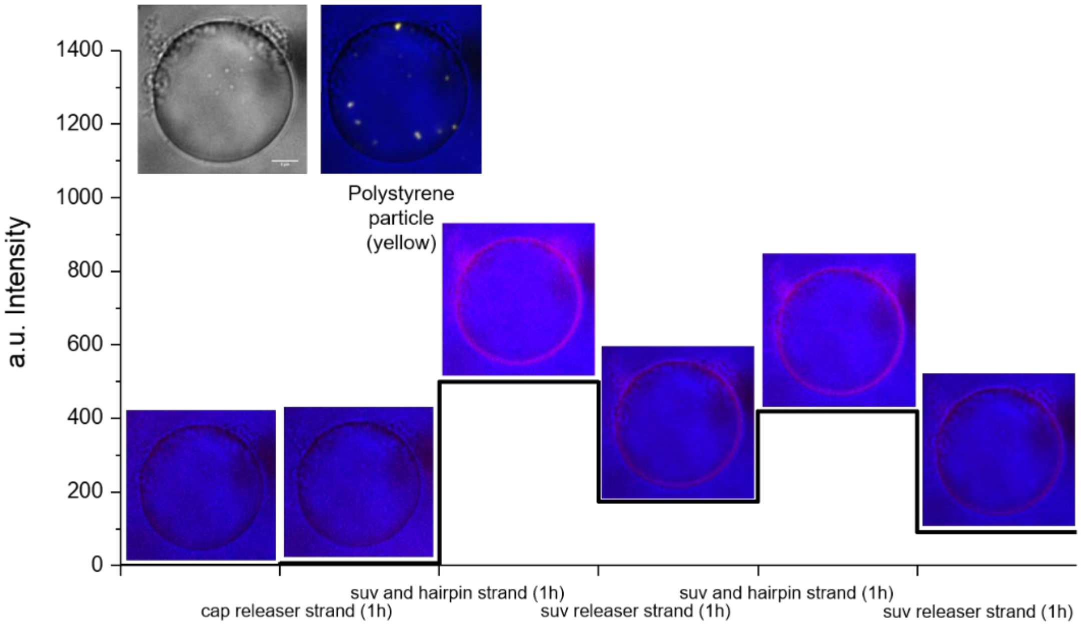


**Figure S7**. Additional reversible aggregation experiment with 2 cycles of adding SUVs, hairpin strands, and SUV releasers. For each step, we kept 1-hour incubation time, because the change of fluorescence intensity gradually stopped after 40 to 50 mins. Similar to Figure 5, yellow dots in the top left image represents polystyrene particles functionalized with Exo III enzymes. Circular fluorescence ring indicates the aggregation of small vesicles on the surface of the giant vesicle.

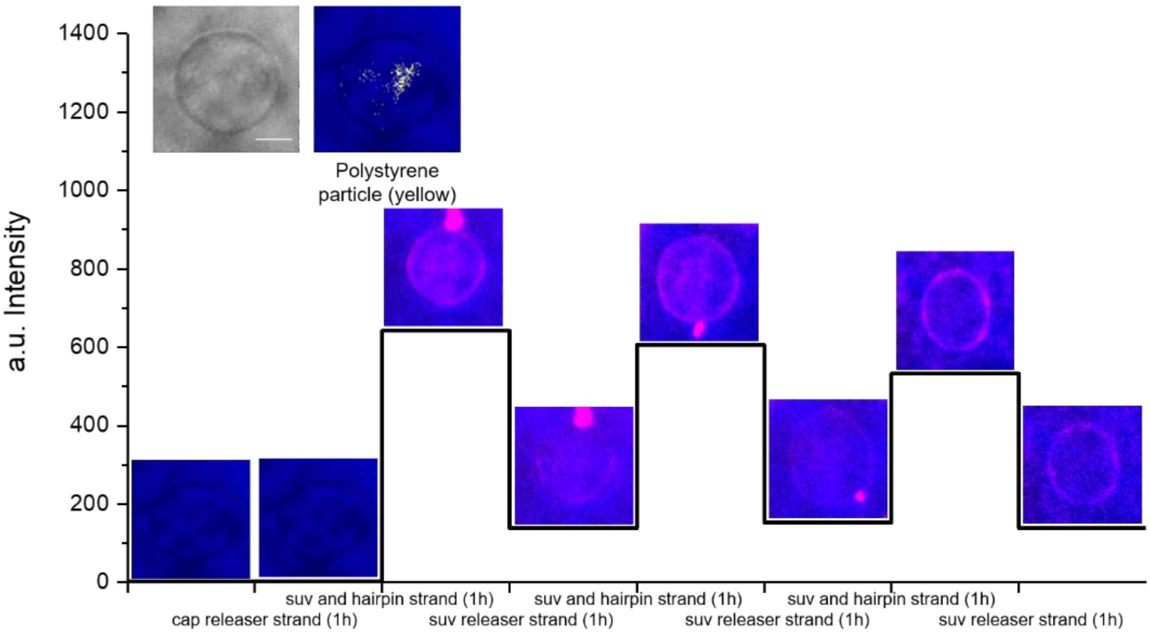

**Figure S8**. Additional aggregation experiment with hairpin signals. Significant changes of fluorescence intensity were observed during the course of experiment. It is noticeable that the GUV immobilized on the glass surface gradually changed its shape from a sphere to ellipsoid. We suspect that this might be caused by the flow in the microfluidic channel.

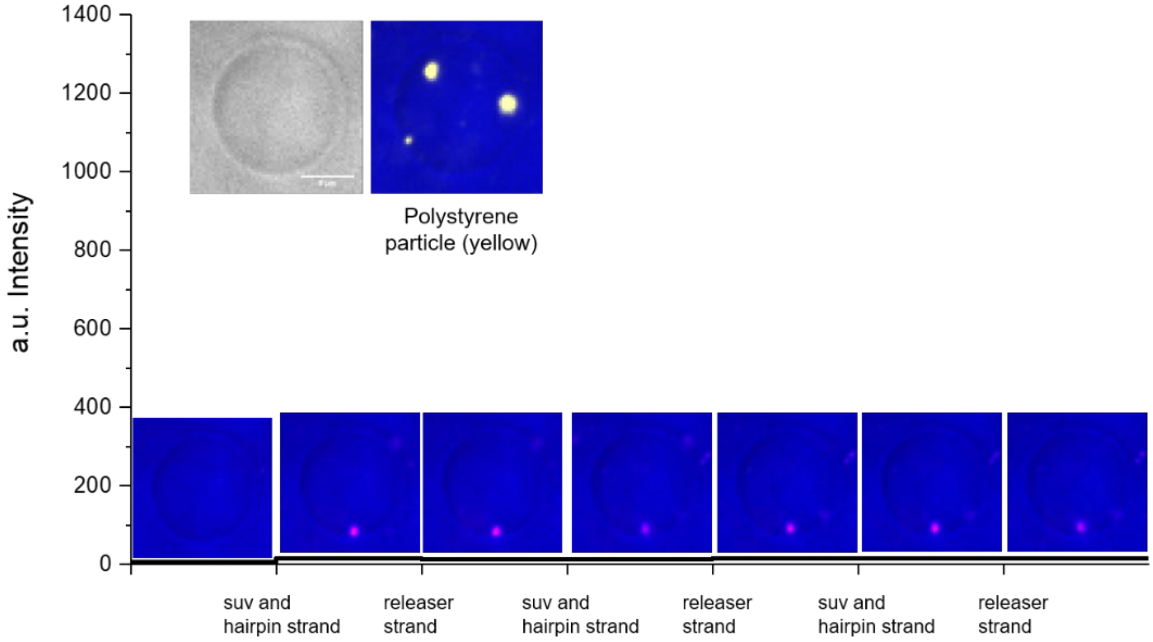


**Figure S9** Control experiment without cap releaser strands. Small vesicles and hairpin strands were introduced into the microfluidic imaging chamber where a giant vesicle was immobilized with DNA pores closed with origami caps. Without adding cap releaser strands, the origami channels on the GUV remained closed; therefore, hairpin strands could not enter the GUV to become active SUV linker strand. As a result, we did not observe significant SUV binding on the GUV. There is only one spot that has nonspecifically bound SUV which was not removed by the SUV releaser strand.

**4. Kinetics**

In Figure 3, we used a single exponential function $I(t) = ae^{k_f t} + b$ for curve-fitting of fluorescence intensity *I(t)* as a function of time *t*, which is related to outflux of GFP and Cy5-DNA molecules from giant vesicles. Here, $k_f$ is the leaking rate, and *b* represents the energy barrier that inhibits further translocation of fluorescent molecules via DNA channels. The outflux kinetics of GFP and Cy5-DNA are expressed as:

$$I_{GFP}(t) = 0.8907e^{-0.3358t} + 0.3692$$
$$I_{DNA\,Dye}(t) = 0.6402e^{-0.1302t} + 0.3321$$

The kinetics of dFITC influx into a vesicle via origami pores in a previous report by Thomsen et al.[7] is expressed as:

$$I_{dFITC}(t) = 0.8106e^{-0.06495t} + 0.1834$$

Thus, the three leaking rates are similar within the order of magnitude ($k_{f,GFP} > k_{f,Cy5\text{-}DNA} > k_{f,dFITC}$) and the time constants are about 3, 8, and 15 mins for GFP, Cy5-DNA, and dFITC, respectively.
